# Innate immune signaling in trophoblast and decidua organoids defines differential antiviral defenses at the maternal-fetal interface

**DOI:** 10.1101/2021.03.29.437467

**Authors:** Liheng Yang, Eleanor C. Semmes, Cristian Ovies, Christina Megli, Sallie Permar, Jennifer B. Gilner, Carolyn B. Coyne

**Author notes:** Address correspondence, Carolyn Coyne, PhD, 3130 Medical Sciences Research Building III 3 Genome Court, Durham, NC 27710 USA.

## Abstract

Infections at the maternal-fetal interface can directly harm the fetus and induce complications that adversely impact pregnancy outcomes. Innate immune signaling by both fetal-derived placental trophoblasts and the maternal decidua must provide antimicrobial defenses at this critical interface without compromising its integrity. Here, we developed matched trophoblast and decidua organoids from human placentas to define the relative contributions of these cells to antiviral defenses at the maternal-fetal interface. We demonstrate that trophoblast and decidua organoids basally secrete distinct immunomodulatory factors, including the constitutive release of the antiviral type III interferon IFN- λ2 from trophoblast organoids, and differentially respond to viral infections through the induction of organoid-specific factors. Lastly, we define the differential susceptibility of trophoblast and decidua organoids to human cytomegalovirus (HCMV) and the transcriptional and immunological responses of these organoids to HCMV infection. Our findings establish matched trophoblast and decidua organoids as *ex vivo* models to study vertically transmitted infections and highlight differences in innate immune signaling by fetal-derived trophoblasts and the maternal decidua.

## Introduction

The maternal-fetal interface is comprised of both the maternal-derived decidua and the fetal-derived placenta, which together support fetal growth, maintain maternal-fetal immunotolerance, and provide immunologic defenses to protect the fetus from infections. The relative contributions of maternal- and fetal-derived cells to these critical functions remain largely undefined given the limitations of models that recapitulate the complex nature of the human maternal-fetal interface.

The fetal-derived placenta contains distinct trophoblast subsets throughout gestation. Trophoblast stem cells located in chorionic villi, the branch like projections that directly contact maternal blood, give rise to cytotrophoblasts (CTBs), the proliferative mononuclear cells of the placenta, the syncytiotrophoblast (STB), a multinucleated contiguous cell layer that covers the villous surfaces, and extravillous trophoblasts (EVTs). EVTs remodel the maternal microvasculature to form spiral arteries, which ultimately deliver maternal blood to the surface of chorionic villi. On the maternal side, decidualization of the endometrium is required for implantation and appropriate placentation. Together, the decidua and placenta maintain immunotolerance throughout gestation and provide the fetus with essential hormones and nutrients to support its growth.

While the placenta and decidua play complementary roles in fetal development, their contributions to antimicrobial defenses are less defined [1, 2]. The placenta forms a formidable barrier to infections, yet certain congenital pathogens (e.g., *Toxoplasma gondii*, Zika virus (ZIKV), HIV, Rubella, and Human Cytomegalovirus (HCMV)) can be vertically transmitted to infect the fetus and in some cases, cause disease and/or pregnancy loss. Studies of tissue explants and histopathologic analysis of the placenta suggest that the decidua and chorionic villi have differing roles in the pathogenesis of infections. The decidua may be a primary site of replication for several microorganisms associated with congenital disease, including HCMV [3-5], ZIKV [6-8], and *Listeria monocytogenes* [9], suggesting it may serve as a reservoir for these infections. By contrast, the placenta is largely resistant to infections and possesses intrinsic mechanisms of innate immune defense. For example, human trophoblasts constitutively release antiviral interferons (IFNs) that restrict infection in both autocrine and paracrine manners [10, 11]. These IFNs are of the type III class, which include IFNs-λ1-3 in humans [12, 13]. Constitutive release of IFNs is a unique feature of trophoblasts as IFNs are tightly regulated in most cell types and induced only after detection of the products of viral replication. Together, these findings suggest that maternal- and fetal-derived tissue differentially respond to and control viral infections; however, the lack of available models to fully interrogate the differences between maternal- and fetal-derived cells has limited a complete understanding of their respective roles in antiviral defense.

The paucity of tractable models to study the human maternal-fetal interface and the limitation of accessing this interface *in vivo* without disrupting pregnancy has limited our understanding of the pathogenesis of and immune responses against many congenital pathogens, including HCMV. HCMV is a species-specific and host-restricted β-herpesvirus that can only infect humans, and animal models using guinea pig (GPCMV) or rhesus macaque (RhCMV) cytomegaloviruses rely on viruses that are genetically disparate from HCMV and do not fully recapitulate the human decidua-placenta interface [14]. Prior studies on placental HCMV infection have used trophoblast progenitor cells, EVT cell lines, and primary trophoblasts; however, these cell lines and isolated primary cells do not model the cellular complexity of the human placenta [15]. Placental and decidual explants have also been employed to model HCMV infection, yet these human explants have high inter-individual variability, which hinders reproducibility, and a limited window of viability in cultured *ex vivo* tissue [4, 6, 16]. Therefore, novel models of the human maternal-fetal interface are needed to understand the contributions of maternal decidual cells and fetal-derived placental cells to immune defense and pathogenesis of placental HCMV infection.

The recent development of trophoblast and decidua organoids from first trimester placental tissue has greatly expanded the models available to study the human maternal-fetal interface [17, 18]. Previous work has shown that isolation and propagation of trophoblast stem cells from first trimester placental chorionic villi leads to the development of three-dimensional organoids that differentiate to contain all lineages of trophoblast cells present in the human placenta and release well-known pregnancy hormones such as human chorionic gonadotropin (hCG) [17]. Similarly, decidua organoids isolated from uterine glands recapitulate the transcriptional profile of their tissue of origin and respond to hormonal cues [18]. These organoid models are ideal to interrogate differences between maternal- and fetal-derived cells; however, decidua and trophoblast organoids have not yet been used to model congenital infections.

To directly compare antiviral signaling pathways between the human placenta and decidua, we generated trophoblast and decidua organoids from matched mid-to-late gestation (26-41 weeks of gestation) human placentas and profiled their release of cytokines, chemokines, and other factors at baseline and under virally infected conditions. These studies revealed that trophoblast organoids basally secrete the IFN-λ2, which does not occur in matched decidua organoids. To determine whether maternal- and fetal-derived cells respond differently to viral infections, we compared the transcriptional profiles and immunological secretomes from organoids treated with a ligand to stimulate toll-like receptor 3 (TLR3) signaling. Trophoblast and decidua organoids differentially responded to TLR3 activation through the secretion of organoid-specific cytokines and chemokines. Lastly, we defined the differential susceptibility of trophoblast and decidua organoids to HCMV infection and identified key differences in the innate immune responses of these organoids to HCMV infection. Together, these studies highlight the differential responses of decidual and placental cells at the maternal-fetal interface to viral infections and provide new models to understand innate immune responses against placental infections.

## Results

### Establishment and characterization of human trophoblast and decidual organoids from mid-to-late gestation tissue

To generate matched human trophoblast organoids (TOs) and decidual organoids (DOs), we used human placental tissue isolated from the second and third trimesters of human pregnancy (26-41 weeks gestation) and cultured progenitor cells isolated from chorionic villi or decidua (**Figure 1A**). Given that previous work developed TOs utilizing tissue isolated from the first trimester [17-22], we optimized the isolation and propagation procedures for isolation of cells from mid-to-late gestation chorionic villi. This optimization included extended mechanical disruption of tissue and the addition of supplements including nicotinamide to promote progenitor cell differentiation. Although the abundance of trophoblast stem/progenitor cells decreases after the first trimester [17], we successfully isolated progenitor cells and generated organoids from all placental tissue harvested (eleven unique placentas). Consistent with the presence of progenitor cells in full-term chorionic villi, we identified cells positive for the proliferation marker Ki67 in villi isolated from two late gestation placental villi used to generate TOs (**Supplemental Figure 1A**). Isolated progenitor cells were embedded in Matrigel to allow for self-organization and propagated for ∼2-3 weeks to generate organoid structures that were morphologically like those previously isolated from first trimester placentas [17, 18] (**Figure 1B, Supplemental Figure 1B**). Once established, organoids could be continuously expanded and passaged every 3-5 days for DOs and 5-7 days for TOs and were highly proliferative based on Ki67 immunostaining (**Supplemental Figure 1C**). Confocal microscopy for cytoskeletal and adhesion markers including cytokeratin-19, actin, E-cadherin, and the Epithelial Cell Adhesion Molecule (EpCAM) confirmed the three-dimensional architecture of established organoids and verified that TOs and DOs expressed markers consistent with their tissues of origin (**Figure 1C, 1D** and **Supplemental Figure 1D**). This also included the expression of the trophoblast-specific marker Sialic Acid Binding Ig Like Lectin 6 (SIGLEC6) in TOs but not DOs and the expression of the mucin MUC5AC in DOs but not TOs (**Figure 1C, 1D** and **Supplemental Figure 1D**). In addition, we confirmed that TOs differentiated to multiple lineages of trophoblast cells and a subset of cells were positive for the EVT specific marker HLA-G (**Figure 1C, Supplemental Figure 1D**). We also found that TOs, but not DOs, secreted the trophoblast-specific hormone hCG as assessed by Luminex assays and OTC test strips (**Figure 1E**). The levels of hCG were stable in TOs propagated for extended periods of time (**Figure 1F**). In addition, we confirmed that DOs released high levels of the secreted mucin MUC16 as determined by Luminex assays, which was not present in TOs (**Figure 1G**). Collectively, these data show that TOs and DOs are morphologically and functionally distinct and that they recapitulate human placental trophoblasts and decidua glandular epithelial cells.

**Figure 1.**
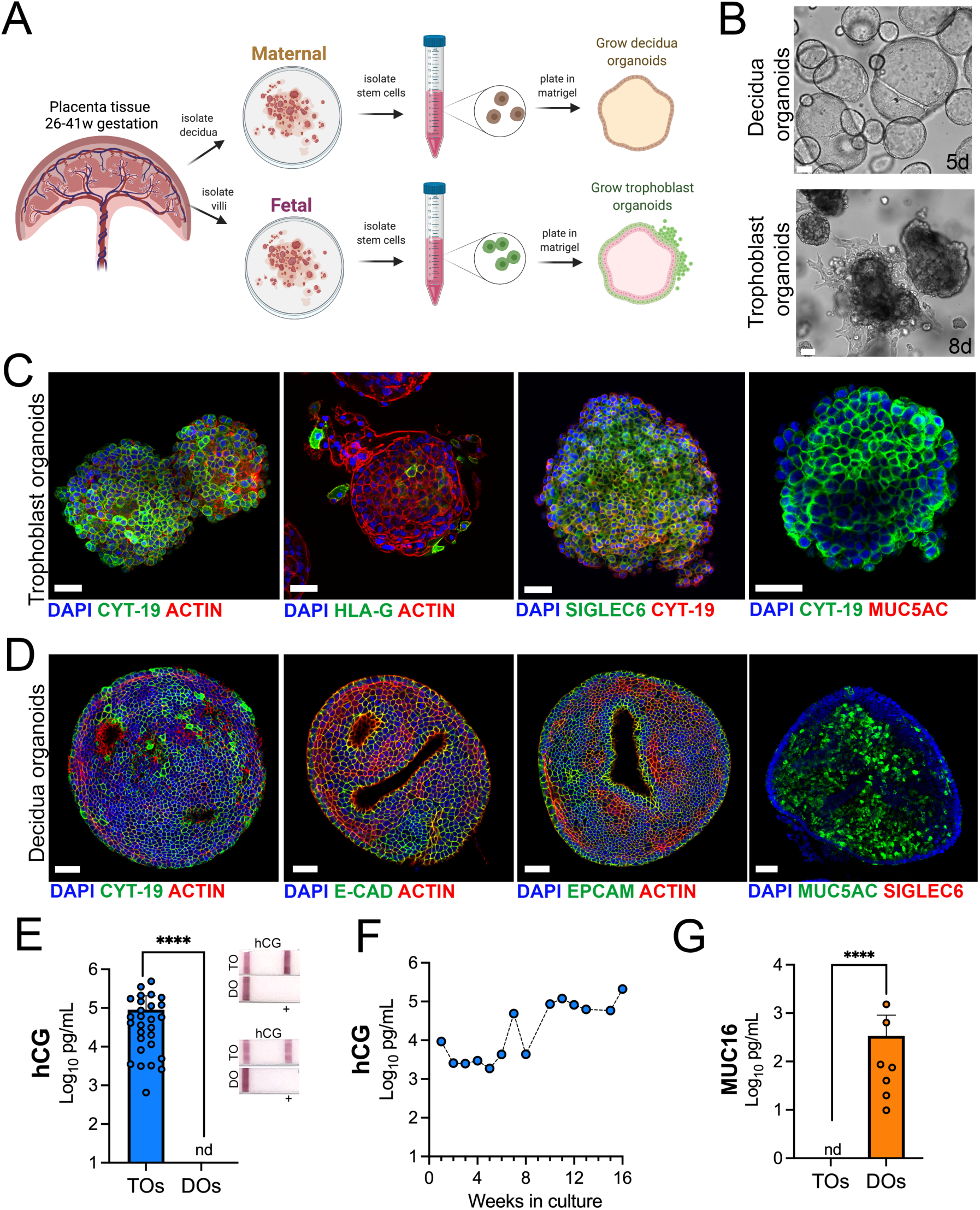
Long-term three-dimensional organoids cultures can be established from human mid- and late-gestation placental tissue. **(A)** Schematic representation of the derivation of trophoblast organoids (TOs) and decidual organoids (DOs) from mid-to-late gestation placental tissue. **(B)** Representative bright-field images of TOs and DOs after passaging in complete growth medium TOM and ExM, respectively at 5 days (DO, top) or 8 days (TO, bottom) post-passaging. Scale bar, 50 µm. **(C)** Confocal microscopy in TOs immunostained for (from left) cytokeratin-19, HLA-G, SIGLEC-6, cytokeratin-19 (in green) and actin, cytokeratin-19, or MUC5AC (in red). DAPI- stained nuclei are shown in blue. Individual channels are shown in Supplemental Figure 1D. Scale bar, 50 µm. **(D)** Confocal microscopy in DOs immunostained for (from left) cytokeratin-19, E-cadherin (E-cad), EpCAM, or MUC5AC (in green) and actin or SIGLEC6 (in red). DAPI-stained nuclei are shown in blue. Individual channels are shown in Supplemental Figure 1D. Scale bar, 50 µm. **(E)** Levels of hCG in conditioned medium (CM) isolated from TOs (blue) or DOs (orange) as determined by Luminex. At right, over the counter (OTC) pregnancy tests for hCG in two matched TO and DO lines. **(F)** Levels of hCG in CM isolated from TOs throughout a 16-week culture period as determined by Luminex. **(G)** Levels of secreted Mucin-16 (MUC16) in CM isolated from TOs (light blue) or DOs (orange) as determined by Luminex assays. In (E) and (G) each symbol represents an individual CM preparation and significance was determined by Mann-Whitney U test. ****, P < 0.0001 and nd, not detected.

### Distinct transcriptional profiles in TOs and DOs as assessed by RNASeq

Next, we performed transcriptional profiling of established TO and DO lines using bulk RNASeq. We found that TOs and DOs were transcriptionally distinct based on differential expression analysis using DESeq2 followed by principal component analysis (PCA) (**Figure 2A**, **Supplemental Figure 2A, 2B, 2C**). TOs highly expressed transcripts specifically enriched in the placenta, including HSD3B1, XAGE2 and 3, GATA3, and ERVW-1, which were absent or expressed at very low levels in DOs (**Figure 2B, Supplemental Figure 2C, 2D, 2E**). In addition, TOs highly expressed transcripts for PSGs, ERVW-1, and CGA, which were not expressed in DOs (**Figure 2B**). The levels of expression of these transcripts were similar to those in primary human trophoblasts (PHTs), confirming that TOs recapitulate the transcriptional signature of primary trophoblasts (**Figure 2B**). These markers were consistent with the presence of the STB (e.g., CGA, PSGs, HSD3B1), CTBs (e.g., TP63), and EVTs (e.g., ITGA5) (**Figure 2B**). In contrast, DOs expressed markers associated with the decidua and decidual stromal cells, including MUC1, MUC5A, MUC5B, SOX17, and HOXB8 which were absent or expressed at low levels in TOs and PHT cells (**Figure 2B, Supplemental Figure 2F, 2G**). To further confirm that TOs and DOs recapitulated their tissues of origin, we profiled the expression of three members of the chromosome 19 microRNA family (C19MC), which are specifically expressed in the human placenta [23]. We found that miR-516B-5p, miR-517A-3p, and miR-525-5p were expressed at high levels in TOs, comparable to levels observed in freshly isolated chorionic villi, but were not expressed in DOs (**Figure 2C**). Lastly, we found that TOs isolated from placentas obtained from male fetuses expressed Y-linked genes, which were absent in female-only derived DOs (**Supplemental Figure 2H**). Collectively, these data establish matched TOs and DOs as model systems of the maternal-fetal interface and demonstrate that they can be isolated from tissue after the first trimester, including from mid-to-late gestation.

**Figure 2.**
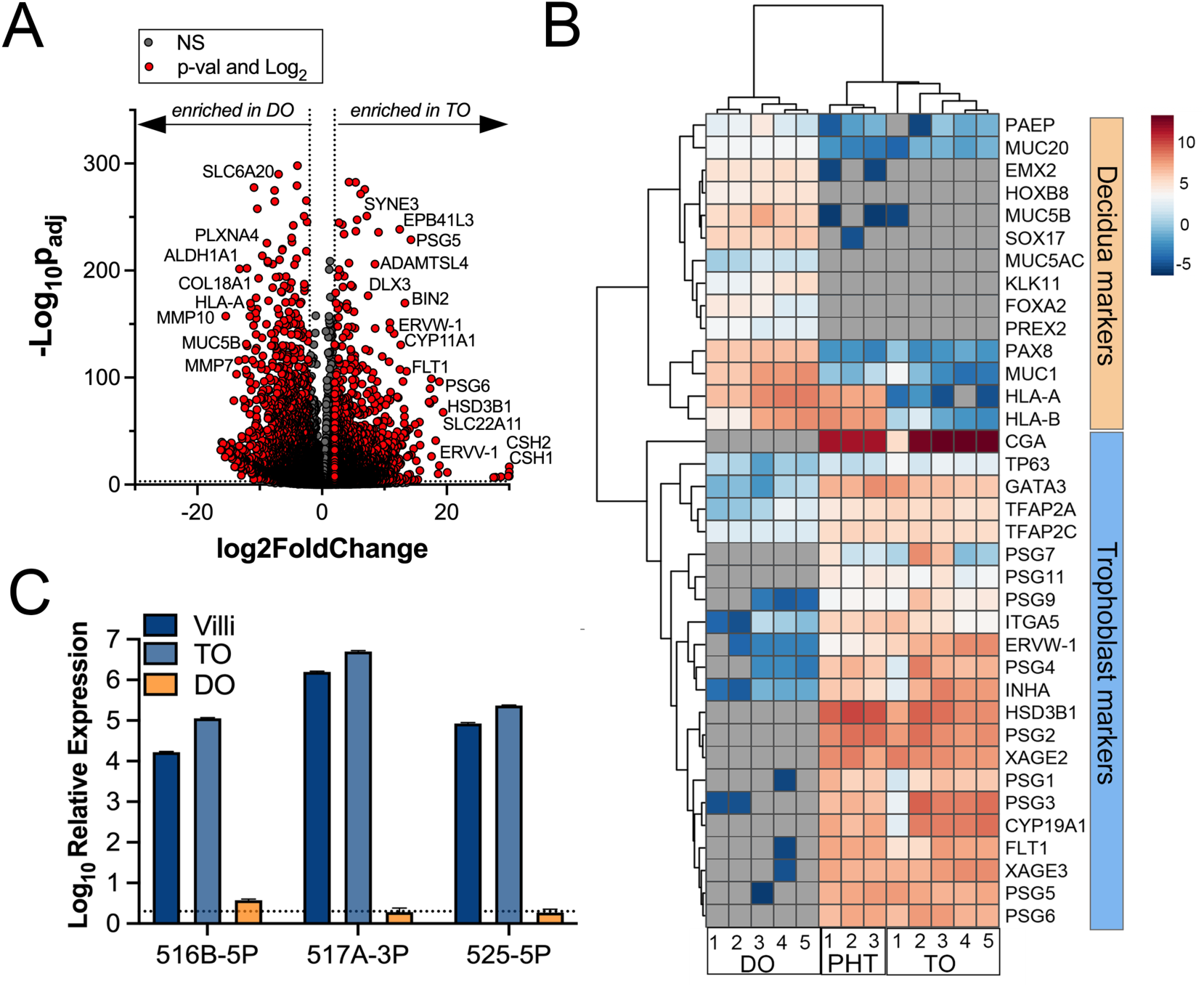
Whole genome transcriptional profiling of trophoblast and decidua organoids. **(A)** Volcano plot demonstrating differentially expressed transcripts in trophoblast organoids (TOs) or decidua organoids (DOs) as assessed by DeSeq2 analysis. Grey circles represent genes whose expression was not significantly changed and red denotes transcripts significantly enriched in TOs (right) or DOs (left) (significance was set at p<0.01 and log2 fold-change of +/- 2). **(B)** Heatmap (based on log2 RPKM values) of transcripts expressed in DOs, TOs, or primary human trophoblasts (PHT) cells that are associated with markers of trophoblasts (top, blue) or the decidua (bottom, orange). Key at right. Red indicates high levels of expression, blue indicates low levels of expression, and grey indicates no reads detected. Hierarchical clustering is shown on left. **(C)** Expression of three members of the C19MC family (miR-516-5p, 517A-3p, and 525-5p) in TOs (light blue) and DOs (orange), or in chorionic villi (dark blue) isolated from human placentas.

### Comparative immune secretome profiling of TOs and DOs

We have shown previously that human chorionic villous explants release numerous cytokines, chemokines, and growth factors that impact both maternal and fetal antimicrobial defenses [11, 24]. Similarly, we have shown that PHTs isolated from mid-gestation and full-term placentas secrete some of these same immunomodulatory factors [11, 24]. We therefore profiled the immunological secretome of matched TOs and DOs isolated from independent placentas to define basal differences in the secretome of cells that comprise the maternal-fetal interface. To do this, we used multianalyte Luminex-based profiling of 105 cytokines, chemokines, growth and other factors from conditioned media (CM) isolated under basal unstimulated conditions. Factors whose secretion was reliably measured over 50pg/mL were defined as present in CM. We found that TOs secreted only two cytokines (IL-6 and IL-28A/IFN-λ2) and three immune-regulated secreted soluble receptors (sTNF- R1, gp130/sIL-6Rβ, sTNF-R2) at baseline (**Figure 3A**). By contrast, DOs released a larger number of immune factors overall, with 21 detected in DO-derived CM at baseline (**Figure 3B**). PCA of these Luminex profiles identified factors contributing to the differential nature of the secretome between TOs and DOs, which included the enrichment of factors including CXCL1, CXCL5, and IL-8 in DOs versus IL-6 and IL28A/IFN-λ2 in TOs (**Figure 3C-F**). Together, these data define the unique immunologic secretome of key cells that comprise the maternal-fetal interface.

**Figure 3.**
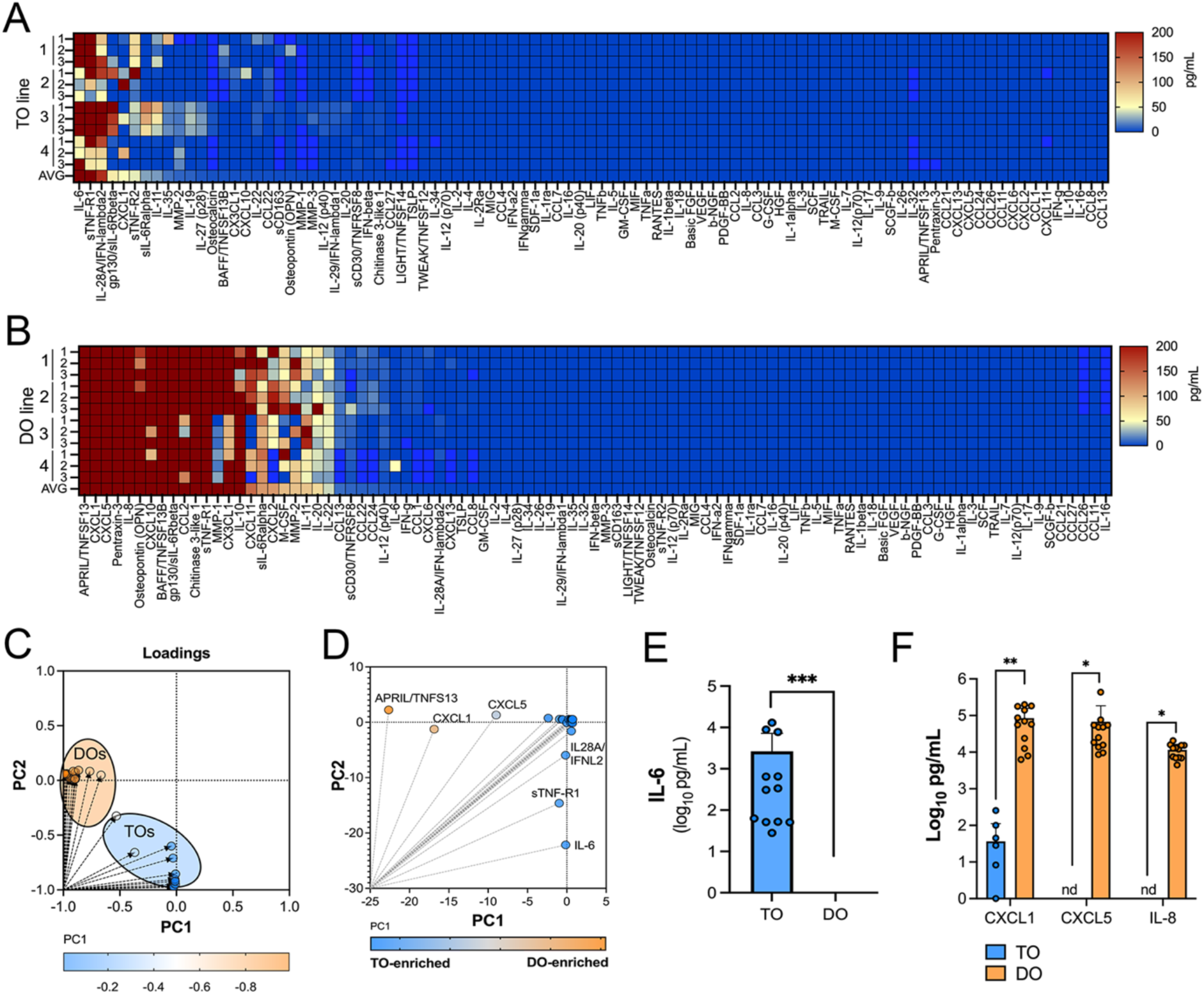
Basal cytokine and chemokine secretion profiles from matched trophoblast and decidua organoids. **(A, B)** Heatmap of twelve conditioned medium (CM) preparations generated from four established TO (A) and DO (B) lines analyzed by Luminex-based multianalyte profiling for the indicated cytokines, chemokines, and growth factors (at bottom). Scale is shown at right. **(C, D)** Principal component analysis of Luminex data shown in panels A and B indicates that the basal secretion between TOs and DOs is unique (C) and identifies the factors that contribute to these differences (D). **(E)** Basal secretion of IL-6 in TOs, but not DOs, as assessed by Luminex. **(F)** Basal secretion of CXCL1, CXCL5, and IL-8 in CM from TOs (in blue) or DOs (in orange). In (E) and (F) each symbol represents an individual CM preparation and significance was determined by Mann- Whitney U test (E) or Two-way Anova with Šídák’s multiple comparisons tests (F). *** P < 0.001, ** P<0.01, * P<0.05.

### Trophoblast organoids constitutively secrete antiviral IFN-λ2

We showed previously that PHT cells isolated from full-term placentas and mid-gestation chorionic villi constitutively release type III IFNs, which protect trophoblasts from viral infections [10, 11]. We found that TOs recapitulated this phenotype and released high levels of IFN-λ2, but not type I IFNs or IFN-λ1 (**Figure 4A**). In contrast, DOs did not basally secrete any detectable IFNs (**Figure 4B**). The levels of released IFN-λ2 were comparable to those from PHT cells and from matched cultured villous explants (**Figure 4C**). Consistent with the specific constitutive release of IFN-λ2 from TOs, the expression of transcripts associated with type III IFNs was high in TOs compared to DOs (**Figure 4D**). We monitored the release of IFN-λ2 over the growth period of TOs and found that it’s release was detectable by ∼10-days post-plating and reached high levels by >14-days in culture, which correlated with similar trends for IL-6 (**Supplemental Figure 3**). We confirmed that CM isolated from TOs, but not DOs, induced interferon stimulated genes (ISGs) in cells treated with this CM using a HEK293 based reporter assay (**Figure 4E**). Consistent with this, we found that CM isolated from TOs was antiviral and reduced ZIKV infection in CM-treated non-placental cells (**Figure 4F**), confirming that TO-derived CM exerts potent antiviral effects on non-placental cells.

**Figure 4.**
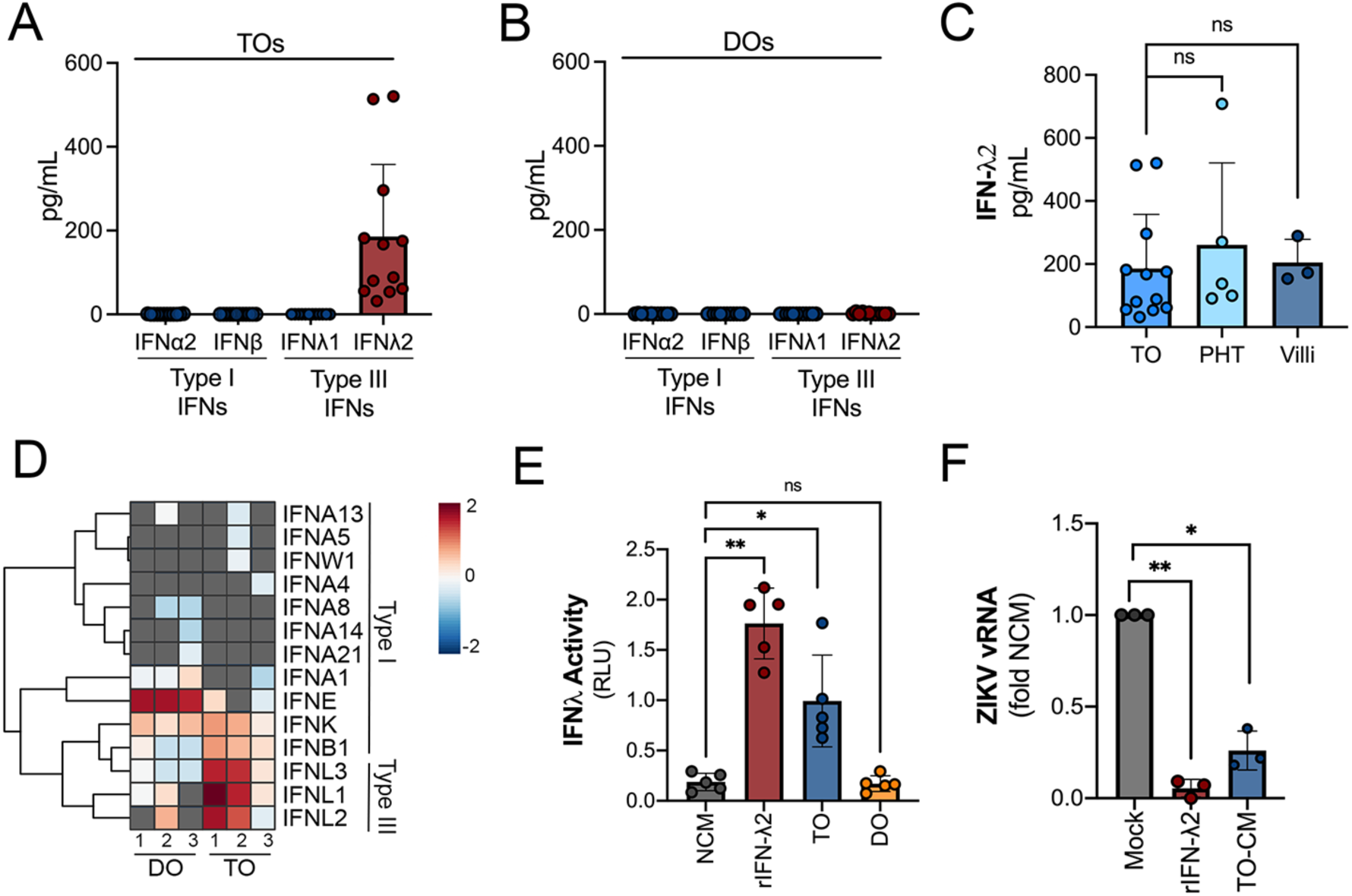
Trophoblast organoids secrete high levels of antiviral IFN-λ2. **(A, B)** Comparison of the levels of the type I IFNs IFN-α2 and IFN-β and type III IFNs IFN-λ1 and IFN-λ2 in conditioned medium (CM) isolated from TOs (A) or DOs (B). **(C)** Comparison of the levels of IFN-λ2 in CM isolated from TOs, matched villous tissue, or in primary human trophoblasts (PHTs). **(D)** Heatmap (based on log2 RPKM values) of transcripts associated with type I (top) or III (bottom) IFNs in TOs or DOs. Key at right. Red indicates high levels of expression, blue indicates low levels of expression, and grey indicates no reads detected. **(E)** IFN-λ activity of CM isolated from TOs (blue) and DOs (orange) as assessed using an IFN-λ reporter 293T cells line. NCM (non-conditioned medium, grey) was as negative control, NCM supplemented with recombinant human IFN-λ2 (100ng) was used as a positive control (red). **(F)** Human osteosarcoma U2OS cells were treated with CM isolated from TOs (blue) or with recombinant IFN-λ2 (100ng) for 24hrs and then infected with ZIKV for 24hrs (in the presence of CM). Level of ZIKV replication was assessed by viral RNA as determined by qPCR. Data are shown as fold change from NCM and from at least three independent CM preparations. Significance was determined by Mann-Whitney U test or by Kruskal Wallace U test with multiple comparisons. *, P < 0.05; **, P < 0.01; ns, not significant. In A-C, E-F, each symbol represents unique CM preparations.

### Differential induction of cytokine and chemokine networks in TOs and DOs in response to poly I:C treatment

Because we observed differences in the basal secretomes of TOs and DOs, we next determined whether TOs and DOs differentially respond to viral infections. To do this, we used poly (I:C), a synthetic ligand of TLR3, and profiled the transcriptional and secretory changes induced by poly I:C treatment of TOs and DOs. We found that both TOs and DOs robustly responded to poly I:C treatment, which corresponded with significant transcriptional changes as assessed by RNASeq (**Figure 5A**, **5B**). Comparison of the transcripts induced by poly I:C treatment (as defined by significance of p<0.01 and log_2_ fold-change of +/- 2) of TOs and DOs revealed that TOs upregulated 448 transcripts whereas DOs upregulated 121 (**Supplemental Figure 4A**). Of these, only 69 were shared between TOs and DOs, the majority of which (64 of 69) were ISGs (**Supplemental Figure 4A, 4B**). Of the differentially induced transcripts, TOs induced the expression of chemokines including CXCL5 (log_2_ fold change 5.9), CX3CL1 (log_2_ fold change 7.1), and CCL22 (log_2_ fold change 7.6), which were the most induced cytokine or chemokine transcripts in poly I:C-treated TOs (**Figure 5C**). In contrast, DOs expressed chemokines at high basal levels, which were not further induced by poly I:C treatment and did not induce any CCL22 (**Figure 5C**). Transcripts most selectively induced in DOs included CD38 and the phospholipase PLA1A (**Figure 5C**).

**Figure 5.**
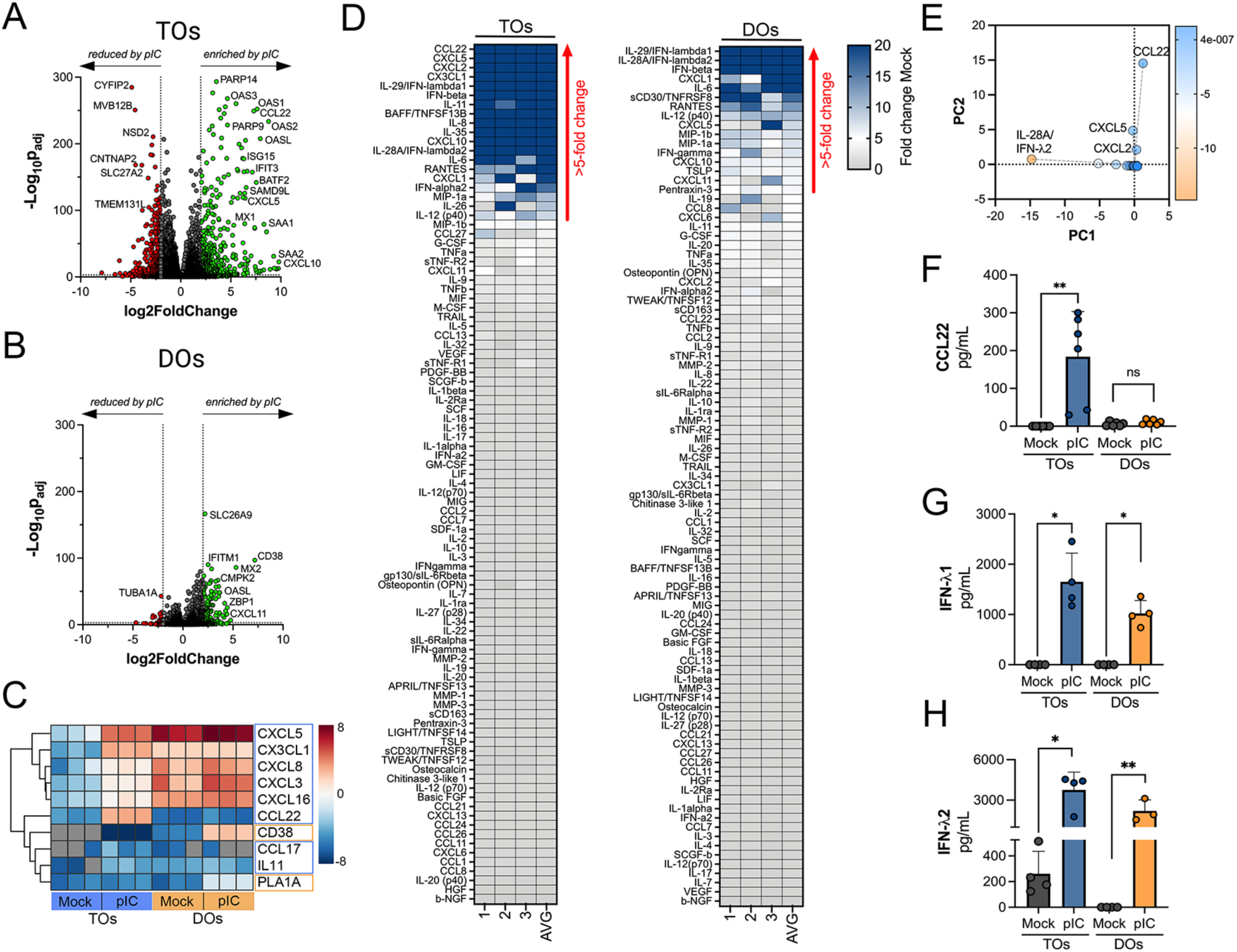
Differential cytokine and chemokine release in TOs and DOs treated with poly I:C. **(A, B)** Volcano plots demonstrating differentially expressed transcripts in trophoblast organoids (TOs, A) or decidua organoids (DOs, B) treated with poly I:C as assessed by DeSeq2 analysis. Grey circles represent genes whose expression was not significantly changed and green denotes transcripts significantly induced by poly I:C treatment whereas red denotes transcripts significantly reduced by poly I:C treatment. Significance was set at p<0.01 and log2 fold-change of +/- 2. **(C)** Heatmap (based on log2 RPKM values) of select transcripts induced uniquely by poly I:C treatment of TOs (blue boxes) or DOs (orange boxes). Key at right. Red indicates high levels of expression, blue indicates low levels of expression, and grey indicates no reads detected. Hierarchical clustering is on the left. **(D)** Heatmaps demonstrating the induction of factors at left (shown as fold change from mock-treated controls) from TOs (left panel) and DOs (right panel) treated with 10μg poly (I:C) and analyzed by multiplex Luminex-based profiling for 105 cytokines, chemokines, and other factors. AVG denotes the average change in concentration of factors over conditioned medium (CM) isolated from three individual preparations. Dark blue denotes significantly induced factors compared with untreated controls. Gray or white denotes little to no change (scale at top right). The red arrow demonstrates factors with induction greater than five-fold change observed in the average of separate experiments. Data are from three individual CM preparations from at least three unique matched organoid lines. **(E)** Principal component analysis of Luminex data shown in left panels identify factors differentially induced by poly I:C treatment of TOs and DOs. **(F-H)** Levels of CCL22 (F), IFN-λ1 (G), and IFN-λ2 (H) in CM isolated from poly I:C-treated TOs (blue) or DOs (orange) or in mock-treated controls (grey). In F-H, each symbol represents individual CM preparations. In F-H significance was determined using a Mann-Whitney U test. **, P < 0.01, *, P < 0.05; ns, not significant.

To confirm the transcriptional changes described above, we performed parallel multianalyte Luminex profiling of 105 cytokines and chemokines in TOs and DOs treated with poly I:C. Both TOs and DOs mounted robust responses to poly I:C treatment that correlated with the release of high levels of select cytokines and chemokines (**Figure 5D**). Cytokines and chemokines with levels >5- fold above baseline were defined as induced by poly I:C treatment. The topmost induced factors in poly I:C-treated TOs were the chemokines CCL22, CXCL5, CXCL2, and CX3CL1 (**Figure 5D**, left), which was consistent with their transcriptional upregulation by RNASeq (**Figure 5C**). PCA of Luminex-based profiling of poly I:C treated TOs and DOs revealed organoid-specific responses to this treatment, which included the specific induction of CCL22 in poly I:C treated TOs (**Supplemental Figure 4C**, **Figure 5E, 5F**). Factors induced by both TOs and DOs included the type III IFNs IFN-λ1 and IFN-λ2 (**Figure 5G, 5H**), which were the top-most induced factors in DOs. In contrast, the type I IFNs IFN-α2 and IFNβ were induced to significantly greater levels in TOs than DOs (**Supplemental Figure 4D, 4E**). Other chemokines induced in TOs were basally expressed at high levels in DOs and were not further induced by poly I:C (**Supplemental Figure 4F-J**).

### Differential susceptibility of TOs and DOs to HCMV infection

HCMV is the most common vertically transmitted pathogen and is associated with high rates of congenital disease, yet how HCMV transmits across the placenta and how the maternal decidua and fetal-derived placenta respond to HCMV infection remains poorly defined. To address these questions, we infected TOs and DOs with HCMV strains AD169r (BADrUL131-Y4, tagged with GFP [25]) and TB40E (tagged with mCherry [26]). We found that DOs were highly permissive to HCMV infection as quantified by detectable GFP or mCherry expression of viral gene expression in the majority of organoids (**Figure 6A, 6C, 6D**). In contrast, TOs were largely resistant to HCMV infection, with low levels of infection by AD169r and almost undetectable levels by TB40E, as assessed by fluorescence of tagged viral gene expression (**Figure 6B, 6C, 6D**). Because we observed minimal HCMV infection in TOs, we next sought to determine whether the low levels of HCMV infection we observed occurred in EVTs, which have been proposed as possible targets of many teratogenic viruses including HCMV [1]. To do this, we quantified the extent of overlap between AD169r infection as assessed by GFP expression and HLA-G, an EVT specific marker. We found that there was almost no overlap between GFP and HLA-G, suggesting that EVTs may not be the primary targets of HCMV infection in TOs (**Figure 6E, 6F**).

**Figure 6.**
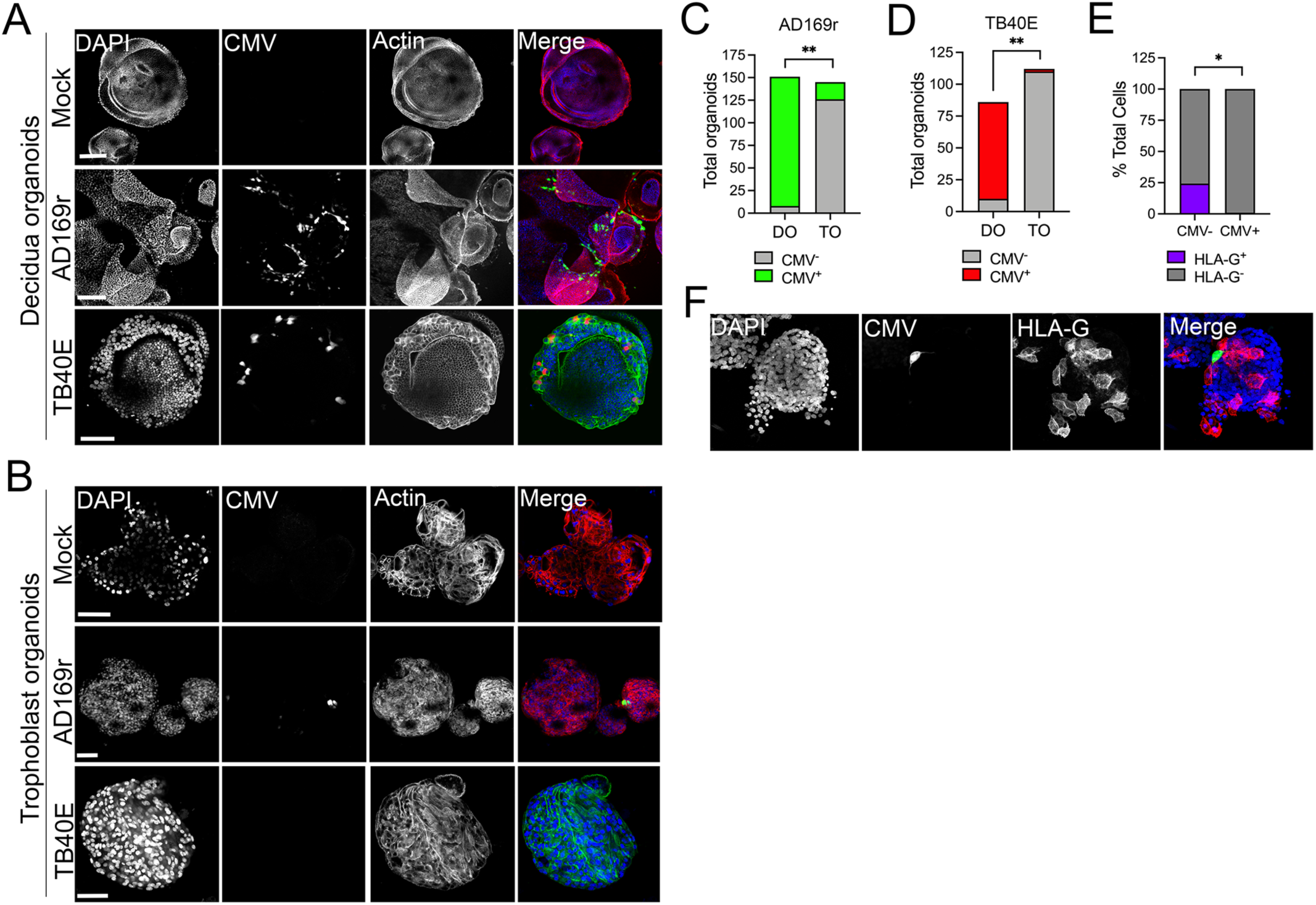
Differential susceptibility of TOs and DOs to HCMV infection. **(A, B)** Representative confocal micrographs of DOs (A) or TOs (B) infected with human CMV (HCMV) strains EGFP-tagged AD169r and mCherry-tagged TB40E for ∼48hrs. Mock infected shown in top row of A and B. Infected cells are positive for EGFP (AD169r, middle row) or mCherry (TB40E, bottom row). Organoids were stained for actin (red in mock- and AD169r-infected or green in TB40E infected). DAPI-stained nuclei are shown in blue. **(C, D)** Quantification of HCMV infection in DOs or TOs infected with AD169r (C) or TB40E (D). Between 75-150 organoids were imaged and HCMV infection scored as at least one positive cell per organoid. Shown are the proportion of organoids that were positive for HCMV. **(E)** Quantification of HCMV (AD169r strain) infection in cells staining positive for HLA-G in TOs. Shown is the percentage of total cells that were positive for HCMV and positive or negative for HLA-G (HLA-G^+^ or HLA-G^-^, respectively). **(F)** Representative confocal micrographs of TOs infected with EGFP-tagged AD169r and co-immunostained for HLA-G (in red). DAPI-stained nuclei are shown in blue. In A-B, scale bar, 50 µm.

### Differential transcriptional responses of TOs and DOs to HCMV infection

Because we observed significant differences between the susceptibility of TOs and DOs to HCMV infection, we next profiled the transcriptional changes induced by HCMV infection in these organoids. Matched TOs and DOs were infected with AD169r or TB40E strains of HCMV and the effects of this infection was assessed by bulk RNASeq transcriptional profiling. TOs responded to HCMV infection through the differential expression of 282 transcripts (AD169r strain) or 173 transcripts (TB40E strain), of which ∼38% were shared between both strains of HCMV (**Figure 7A, 7C**). By contrast, DOs responded to HCMV infection less robustly, with 95 (AD169r strain) or 117 (TB40E strain) transcripts differentially expressed by this infection (**Figure 7B, 7D**). Of these differentially expressed transcripts, only 3-4% were shared between TOs and DOs infected with AD169r (**Figure 7E**) or TB40E (**Figure 7F**). Of these shared transcripts, only three were induced by both strains of HCMV in both TOs and DOs and included CCNA1 (Cyclin A1), the ISG OAS2, and EPSTI1 (Epithelial Stromal Interaction 1) (**Figure 7G**). Transcripts specifically induced by HCMV infection of TOs included H3Y1 (H3.Y Histone 1), KHDC1L (KH Domain Containing 1 Like), several members of the PRAME Family (PRAMEF2, 8, 12, 14, and 20), and TRIM49A and B (Tripartite Motif Containing 43) (**Figure 7H**). In contrast, DO-specific transcripts included several components of the nitric oxide signaling pathway including NOS2 (Nitric Oxide Synthase 2), DUOX2 (Dual Oxidase 2), and DUOXA2 (Dual Oxidase Maturation Factor 2) as well as CXCL10 and CSF3/G-CSF (**Figure 7I**).

**Figure 7.**
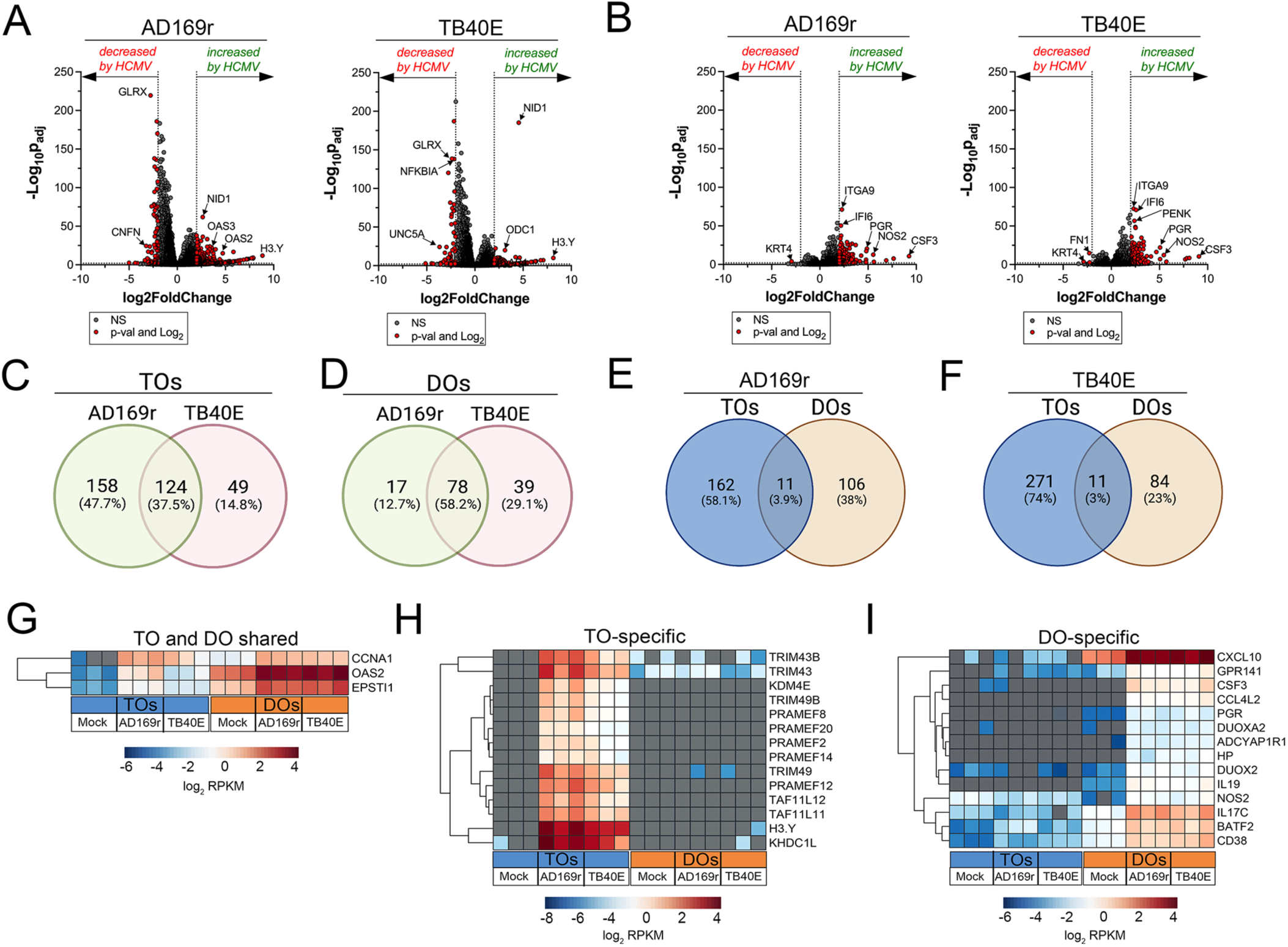
Transcriptional profiling of HCMV infected TOs and DOs. **(A, B)** Volcano plots demonstrating differentially expressed transcripts in trophoblast organoids (TOs, A) or decidua organoids (DOs, B) infected with HCMV strains AD169r or TB40E (as denoted at top) by DeSeq2 analysis. Grey circles represent genes whose expression was not significantly changed and red denotes transcripts significantly changed by HCMV infections. Significance was set at p<0.01 and log2 fold-change of +/- 2. **(C, D)** Venn diagrams of differentially expressed transcripts in TOs (C) or DOs (D) infected with HCMV AD169r (green circle) or TB40E (red circle) strains. **(E, F)** Venn diagrams of the extent of overlap in differentially expressed transcripts in TOs (blue circle) or DOs (orange circles) infected with HCMV AD169r (E) or TB40E (F) strains. **(G)** Heatmap (based on log2 RPKM values) of transcripts induced by HCMV infection of both TOs (blue box, bottom) or DOs (orange box, bottom). **(H, I)** Heatmap (based on log2 RPKM values) of transcripts induced by HCMV infection specifically in TOs (H) or DOs (I). In G-I, key at bottom and red indicates high levels of expression, blue indicates low levels of expression, and grey indicates no reads detected. Hierarchical clustering is on the left.

### Differential immune secretome of TOs and DOs infected with HCMV

To define the innate immune responses to HCMV infections in TOs and DOs, we used Luminex multianalyte assays of >100 cytokines and chemokines in TOs and DOs infected with HCMV strains AD169r or TB40E. We found that TOs responded to HCMV infection by the secretion of relatively few cytokines, which included pentraxin-3 (PTX3), first described as an endothelial factor induced by IL-1β treatment [27], IL-8, and IL-11 (**Figure 8A, 8C, 8D, Supplemental Figure 5A**). In contrast, DOs responded to HCMV infection through the secretion of both proinflammatory cytokines and chemokines (e.g., CXCL10, MIP-1α, MIP-1β, IL-6) and through the release of the type III IFN IFN- λ2 (**Figure 8B, 8C, 8E-F, Supplemental Figure 5B**). In contrast to IFN-λ2, DOs did not induce any other detectable IFNs such as IFN-α2 or IFN-β or IFN- λ1 (**Figure 8B**, **8G-I**). Together, these findings highlight the distinct immunoregulatory and transcriptional cascades induced by HCMV infection of decidual versus placental cells that comprise the human maternal-fetal interface.

**Figure 8.**
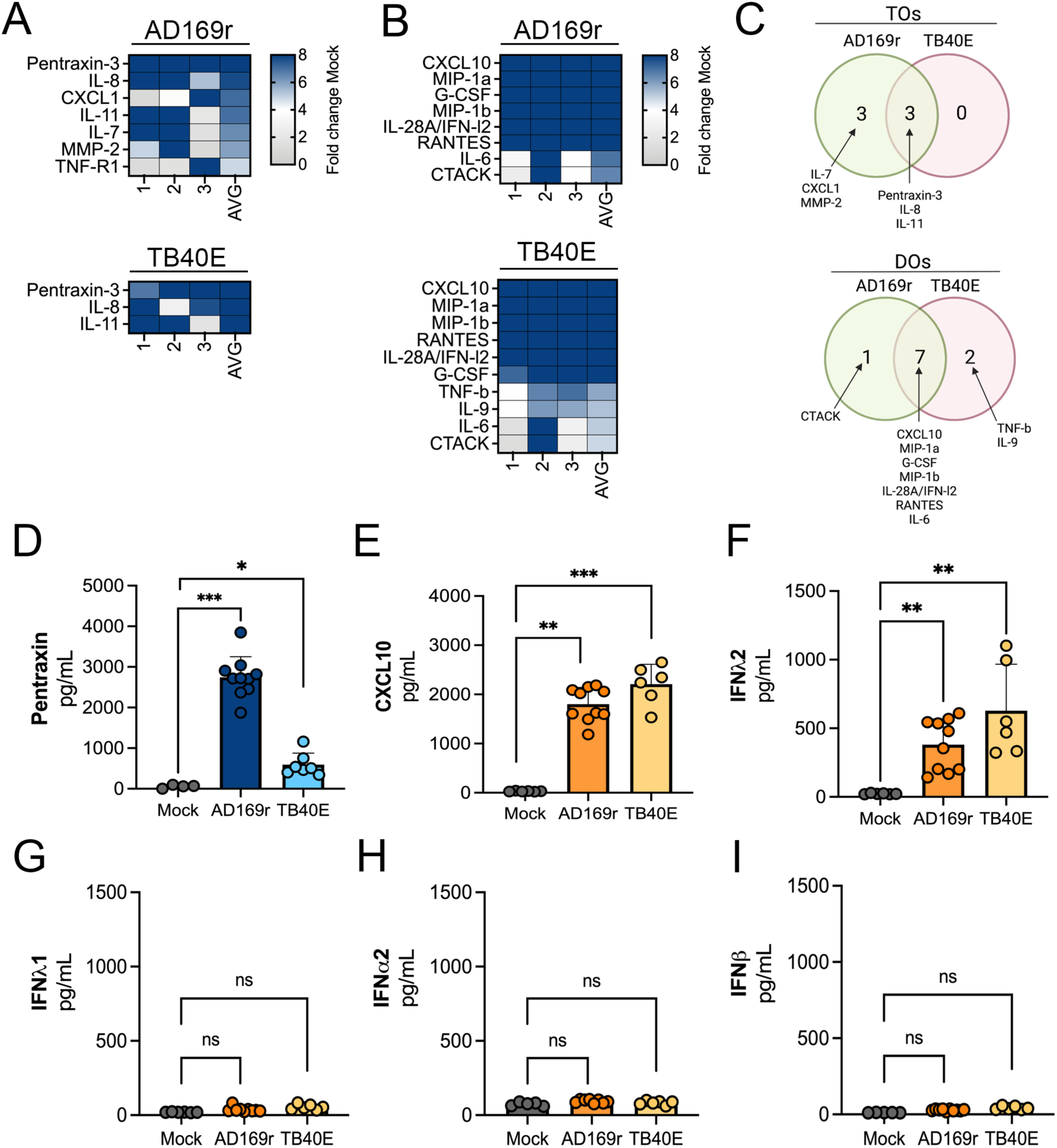
Differential innate immune responses to HCMV infection of TOs and DOs. **(A, B)** Heatmaps demonstrating the induction of factors at left (shown as fold change from mock-treated controls) from TOs (A) and DOs (B) infected with HCMV (AD169r, top or TB40E, bottom strains). Shown are factors induced >5-fold (full heatmap is shown in Supplemental Figure 5). AVG denotes the average change in concentration of factors over conditioned medium (CM) isolated from three individual preparations. Dark blue denotes significantly induced factors compared with untreated controls. Gray or white denotes little to no change (scale at top right). **(C)** Venn diagrams comparing the factors induced by infection of TOs (top) or DOs (bottom) with AD169r (green) or TB40E (red). **(D)** Levels of pentraxin in TOs infected with AD169r (middle, dark blue) or TB40E (right, light blue) compared to mock-infected controls (left, grey). **(E-I)** Levels of CXCL10 (E), IFN-λ2 (F), IFN-λ1 (G), IFN-α2 (H), or IFN-β (I) in CM isolated from DOs infected with HCMV AD169r (middle, dark orange) or TB40E (right, light orange) or in in mock-treated controls (left, grey). In D-I, each symbol represents individual CM preparations and significance was determined using a Mann-Whitney U test. *** P < 0.001, ** P < 0.01, * P < 0.05, ns, not significant.

## Discussion

Our work establishes fetal- and maternal-derived organoids as models for defining innate immune signaling at the maternal-fetal interface. Similar to PHTs [10] and mid-gestation chorionic villi [11], we show that TOs constitutively secrete type III antiviral IFN-λ, that act in paracrine to restrict viral infections. In contrast, although DOs did not basally secrete IFNs, they are immunologically more active under resting conditions and basally secrete cytokines and chemokines not detectable in TOs. In addition, we show that TOs and DOs respond to TLR3 activation via the induction of distinct immune regulatory networks, which includes the specific release of select chemokines such as CCL22 in TOs. Lastly, we used matched TOs and DOs to explore differences in susceptibility to HCMV and to define the differential responses of the maternal decidua and fetal-derived organoids to HCMV infection. Taken together, our work suggests that cells that comprise the maternal-fetal interface defend from viral infections in a cell-type specific manner and further highlight the unique contributions of fetal-derived trophoblasts in antiviral immunity.

The previous establishment of trophoblast and decidua-derived organoids has significantly expanded the *in vitro* systems that can model the complexity of the maternal-fetal interface [17, 18]. These previous studies utilized tissue isolated from the first trimester of pregnancy (6-9 weeks gestation for TOs and 8-12 weeks gestation for DOs) [17, 18], although another study successfully established DOs from full-term tissue [22]. Two-dimensional cultures of stem cell-derived primary trophoblasts were also obtained from the first trimester [20]. The placenta is a dynamic organ that changes throughout gestation. Here, we show that TOs can also be generated from later stages of gestation, including from progenitor cells isolated in the second and third trimesters of pregnancy and full-term placental tissue, which required optimization from conditions required for first trimester tissue. Although the abundance of progenitor cells, as defined by the presence of Ki67 positive nuclei, declines throughout gestation, we show that full-term chorionic villi contain Ki67 positive cells at low levels and that these cells can be isolated to generate organoids. Given that obtaining first trimester tissue can be complicated by access and/or regulatory restrictions, our study provides an additional source for the development of placental organoids from tissue obtained later in gestation. In addition, given that many congenital infections occur in the second and third trimesters of pregnancy, organoids isolated from later stages of pregnancy may better reflect clinically relevant models of congenital infections. For instance, the risk of transplacental transmission of HCMV infection from mother-to-fetus is only 20% in the first trimester but can reach up to 75% in the third trimester, highlighting the relevance of our organoid model for studying placental HCMV infection across gestation [28].

Unlike many other barrier cell types such as the epithelium, human trophoblasts have the unique capacity to robustly secrete cytokines and other immunological factors without stimulation. PHTs from full-term placentas and mid-gestation chorionic villi secrete cytokines that mediate innate immune defenses against bacteria, parasites, and viruses [10, 24, 29]. Consistent with this concept, serum isolated from pregnant women often contains elevated levels of these cytokines, indicating that at least some have reached the systemic circulation [24, 30-32]. Similarly, the decidua releases specific cytokines and alterations in these profiles may be associated with pregnancy complications including miscarriage [33-38]. The decidua also responds to microbial infections via the release of additional cytokines that function in innate defense [7, 39, 40]. However, in many cases, these studies were performed on tissue explants that were multicellular in nature, thus preventing an assessment of the contributions of cytokine release from distinct cell types. Similarly, previous work utilizing tissue explants that were treated with recombinant cytokines, synthetic ligands of various innate immune pathways, or infected with pathogens are also unable to define the contributions of distinct cell types in these responses. The use of organoids allows for analyses of cellular responses to infection or innate immune activation from trophoblasts and decidual glands in primary cell models that contain isolated cell types enriched at the maternal-fetal interface. An additional strength of the TO model is that trophoblast stem cells can differentiate into all three distinct trophoblast cell types comprising the human placenta, which is not possible in many other existing models including isolation of CTBs from full-term placentas, which do not differentiate to contain EVTs.

The constitutive release of IFN-λs from human trophoblasts is a unique phenomenon that runs counter to much of what is known about antiviral IFN signaling, which is centered on a requirement for viral sensing as a prerequisite to induce this pathway. Similar to our previous studies in PHT cells and mid-gestation chorionic villous explants [10, 11], our data show that progenitor cell-derived TOs also constitutively release IFN-λs. Our studies also show that type III IFNs are further induced by activation of antiviral innate immune signaling in both TOs and DOs, with IFN-λs being amongst the most induced cytokines detected in both organoid types. Although IFNs are released from TOs constitutively and upon induction of antiviral signaling, our studies also point to select chemokines as key components in the responses of the human placenta to viral infections. We show that the regulatory T cell (Treg) chemoattractant CCL22 is selectively released from TOs in response to TLR3 activation. We have shown previously that human PHTs and mid-gestation chorionic villi also respond to *T. gondii* infection through the selective release of CCL22 [29, 41], suggesting that this chemokine may be key mediator of the response of the human placenta to microbial infections. Given that Treg infiltration into the decidua has been associated with adverse pregnancy outcomes including miscarriage [42], CCL22 induction may directly impact maternal-fetal tolerance and adversely impact fetal health.

Understanding congenital HCMV transmission and innate immune defenses against HCMV infection at the maternal-fetal interface has been greatly hindered by a lack of model systems and has relied heavily on first trimester explant tissues to-date. Our work highlights that trophoblast and decidua organoids derived from mid-to-late gestation tissue can be leveraged as an accessible model system for studying HCMV infection at the maternal-fetal interface. Our finding that DOs were highly susceptible to HCMV infection is consistent with prior work in first trimester decidua explants demonstrating that HCMV productively infects the maternal decidua, which may function as a viral reservoir for HCMV [4, 43]. Using trophoblast and decidua organoids, we found that DOs had heightened susceptibility to HCMV infection compared to matched TOs. This finding of elevated decidual susceptibility to HCMV has important implications, as an improved understanding of the pathogenesis of viral transmission *in utero* will be key to developing effective interventions to prevent congenital infection. Our transcriptional and Luminex-based profiling also revealed that HCMV-infected TOs and DOs differentially respond to this infection through organoid-specific transcriptional signatures and cytokine and chemokine signaling networks. For example, CXCL10 was highly upregulated in HCMV-infected DOs by both RNASeq and Luminex-based profiling, which is consistent with prior findings of elevated CXCL10 in the amniotic fluid of HCMV-transmitting women [44] and in *ex vivo* infections of decidua explants [6, 43]. Moreover, the induction of IL-8 in HCMV-infected TOs mirrors the IL-8 secretion observed previously in response to HCMV infection of isolated first trimester trophoblasts [45] and placental explants [6]. That our *in vitro* organoid infections recapitulated innate immune responses previously observed *in vivo* and *ex vivo* further validates our model of matched decidua and trophoblast organoids for studying HCMV and other vertically transmitted pathogens.

Our work presented here develops TOs and DOs from mid-to-late gestation placental tissue and demonstrates that these organoids can be used to study mechanisms of microbial vertical transmission and antiviral innate immune signaling. We show that TOs and DOs constitutively secrete unique cytokines and chemokines and respond to viral infections through the release of organoid-specific immunomodulatory factors. Our work also highlights the prominent role of type III IFNs in antiviral immunity at the maternal-fetal interface and demonstrates that TOs constitutively release IFN-λs and that DOs can be induced to release IFN-λs following viral infection. Collectively, these studies define the differential responses of decidual and placental cells comprising the maternal-fetal interface to viral infections and suggest that organoid-based models can be used to define innate immune signaling at this interface.

## Materials and Methods

### Human samples

Human tissue used in this study was obtained through the UPMC Magee-Womens Hospital Obstetric Maternal & Infant Database and Biobank or from Duke University after approval was received from the University of Pittsburgh or Duke University Institutional Review Board (IRB) and in accordance with the guidelines of the University of Pittsburgh and Duke University human tissue procurement. Placental tissue used in this study was collected from the second (23^rd^-27^th^ weeks) and third (35^th^- 41^st^ weeks) trimesters. Placental tissue was excluded for diagnosis of PPROM or maternal infection.

### Derivation and culture of trophoblast organoids (TO) from human placental samples

Villous trophoblast stem/progenitor cells were isolated from fresh placental tissues as described previously with further optimization based on the characteristics of late gestation placenta villi [17]. Briefly, villi collected from fresh placental tissue were intensively washed, then were sequentially digested with 0.2% trypsin-250 (Alfa Aesar, J63993-09)/0.02% EDTA (Sigma-Aldrich, E9884-100G) and 1.0 mg/mL collagenase V (Sigma-Aldrich, C9263-100MG) and further mechanically disrupted by pipetting up and down vigorously ∼10 times with 10 ml serological pipette. Pooled digests were washed with Advanced DMEM/F12 medium (Gibco 12634-010) and pelleted by centrifugation, then resuspended in approximate 10 times volume of ice-cold growth-factor-reduced Matrigel (Corning 356231). Matrigel “domes” (40 µl/well) were plated into 24-well tissue culture plates (Corning 3526), placed in a 37°C incubator to pre-polymerize for ∼3 min, turned upside down to ensure equal distribution of the isolated cells in domes for another 10 min, then carefully overlaid with 500 µL prewarmed trophoblast organoid medium (TOM) consisting of Advanced DMEM/F12 (Life Technologies, 12634-010) supplemented with 1× B27 (Life Technologies, 17504-044), 1× N2 (Life Technologies, 17502-048), 10% FBS (vol/vol, Cytiva HyClone, SH30070.03), 2 mM GlutaMAX^TM^ supplement (Life Technologies, 35050-061), 100 µg/mL Primocin (InvivoGen, ant-pm-1), 1.25 mM N-Acetyl-L-cysteine (Sigma, A9165), 500 nM A83-01 (Tocris, 2939), 1.5 µM CHIR99021 (Tocris, 4423), 50 ng/mL recombinant human EGF (Gibco, PHG0314), 80 ng/mL recombinant human R-spondin 1 (R & D systems, 4645-RS-100), 100 ng/mL recombinant human FGF2 (Peprotech, 100- 18C), 50 ng/mL recombinant human HGF (Peprotech, 100-39), 10mM nicotinamide (Sigma, N0636- 100G), 5 µM Y-27632 (Sigma, Y0503-1MG), and 2.5 µM prostaglandin E2 (PGE2, R & D systems, 22-961-0). Cultures were maintained in a 37°C humidified incubator with 5% CO2. Medium was renewed every 2-3 days. Small trophoblast organoid spheroids became visible by approximately day 12 post-isolation. Derived TOs were passaged every 5 to 7 days depending on their size and density. To passage, TOs were released from domes with ice-cold cell recovery solution (Corning, 354253) and then were digested in prewarmed TrypLE Express (Gibco, 12605-028) or Stem Pro Accutase (Gibco, A11105-01) at 37°C for 5-7 min. Further mechanical dissociation was achieved with an Eppendorf Explorer Plus electronic pipettes at its “mix” function setting by pipetting up and down around 400 times through small-bore pipette tips. Dissociated TOs fragment were collected and washed by centrifuge, then resuspended in fresh ice-cold Matrigel and replated as domes at the desired density for continuous culture. For freezing TOs, overlaid TOM was aspirated, TOs with Matrigel were resuspended in CryoStor CS10 stem cell freezing medium (STEMCell Technologies, 07930) frozen at -80 ℃ and then transferred to liquid nitrogen for long-term storage. For thawing cryopreserved TOs, organoids were thawed as quickly as possible, diluted with 5 times volume of basic TOM containing Advanced DMEM/F12, 2 mM GlutaMAX supplement, 10 mM HEPES (Gibco, 15630-106), 1 × Penicillin/Streptomycin (Lonza, 17-602E) and centrifuged to pellet. Afterwards, TOs were resuspended in new ice-cold Matrigel and replated for recovery culture and passaged as described below.

### Derivation and culture of decidual organoids (DOs) cultures from human decidua samples

Decidual gland-enriched cell suspension were acquired from fresh decidual tissue as described previously [18, 46]. Briefly, isolated decidual tissues were cut into small pieces then digested in 1.25 U/mL Dispase II (Sigma, D4693)/0.4 mg/mL collagenase V (Sigma, C-9263) solution. Glandular cells were filtered out from the digestion solution supernatant and resuspended in ice-cold Matrigel (Corning, 356231), plated as domes in 24-well plates (Corning 3526), and finally overlaid with 500 µL prewarmed organoid Expansion Medium (ExM) with the same composition as previously described[18]. The growth medium was renewed every 2-3 days. Mature organoids were passaged by mechanically disruption following TrypLE Express (Gibco, 12605) digestion every 3-5 days. Other maintenance were similar to those as previously described [46].

### Generation of TO and DO conditioned media (CM)

Overlaid TOM and ExM were collected every 2-4 days based on the growth status of TOs and DOs. Media from at least ten timepoints for each established organoid lines were collected as TO-CM or DO-CM. Non-conditioned medium (NCM) was TOM or ExM (described above) that had not been exposed to TOs or DOs, respectively.

### Cells and Viral Infections

Human osteosarcoma U2OS (ATCC HTB-96) and Vero (ATCC CCL-81) cells were grown in Dulbecco’s minimal essential medium (DMEM) with 10% FBS and 1% antibiotic. Human foreskin fibroblast (HFF), JEG-3 and JAR were purchased from the ATCC and cultured as previously described [10, 47]. HEK-Blue IFN-λ reporter cells (hkb-ifnl) were purchased from InvivoGen and cultured according to the manufacturer’s instructions. ZIKV (Paraiba/2015, provided by David Watkins, University of Miami) was propagated in Vero cells. Viral titers were determined by fluorescent focus assay as previously described [48] using recombinant anti-double-stranded RNA monoclonal antibody J2 (provided by Abraham Brass, University of Massachusetts) in Vero cells. Infection was determined by qRT-PCR as described below. To assess antiviral activity, U2OS cells were exposed to TO or DO CM, or matched NCM, 24 hr prior to infection with ZIKV Paraiba/2015 for 72hrs. GFP-tagged AD169r (BADrUL131-Y4, a gift from T. Shenk [25]) and mCherry-tagged TB40/E (a gift from N. Moorman [26]) were propagated by infecting HFF cells (MOI = 0.01) followed by incubation for ∼14 days until 90-95% of cells showed cytopathic effect. Cells and supernatant were harvested, filtered (0.45-μm), and then concentrated on a 20% sucrose cushion (20,000 rpm) before the virus pellet was resuspended in sterile media before titration. HCMV infection of TOs and DOs was performed using 3-4×10^6^ PFU virus per well for 48hrs. For infections, TOs and DOs were plated directly onto 15μL of a Matrigel coating in each well of an 8-well chamber slide (Millicell, EZslide, Millipore) or onto 20μL of Matrigel coating in 24-well plates and cultured for 5-7 days prior to infection. After 48hrs, conditioned media was harvested and stored at -80 °C until Luminex profiling and cells were fixed with paraformaldehyde for 15mins before staining for immunofluorescence microscopy, as detailed below.

### Immunofluorescence microscopy

TOs and DOs were released from Matrigel domes with 1 mL of cell recovery solution (Corning, 354253) per well without disrupting their 3D architecture, then were fixed in 4%PFA for 30 min at room temperature, followed by 0.5% Triton X-100/PBS to permeabilize for 30 min at 4°C. Organoids were washed and blocked in 5%(v/v) goat serum/0.1%(v/v) Tween-20 in PBS for 15 min at room temperature and then incubated with primary antibodies in the above-described blocking solution at 4°C overnight. Organoids were then washed with PBS and then incubated for 1-2 h at room temperature with Alexa Fluor-conjugated secondary antibodies (Invitrogen). Organoids were washed again with PBS and mounted in Vectashield (Vector Laboratories) containing 4′,6-diamidino- 2-phenylindole (DAPI) and were transferred onto microscope slides with large-orifice 200 µL-tips (Fisher Scientific, 02707134). The following antibodies or reagents were used: cytokeratin-19 (Abcam, ab9221), E-cadherin (Invitrogen, PA5- 85088), EpCAM (Abcam, ab223582), Mucin 5AC (Abcam, ab3649), SIGLEC6 (Abcam, ab262851), HLA-G (Abcam, ab52454), Ki67 (550609, BD Biosciences), Alexa Fluor 594–conjugated phalloidin (Invitrogen, A12381), Alexa Fluor Plus 488 Goat anti-Mouse IgG secondary antibody (Invitrogen, A32723), Alexa Fluor 594 Goat anti-Mouse IgG secondary antibody (Invitrogen, A11032), Alexa Fluor 488 Goat anti-Rabbit IgG secondary antibody (Invitrogen, A11034), Alexa Fluor 594 Goat anti-Rabbit IgG secondary antibody (Invitrogen, A11037). Images were captured using Zeiss 880 Airyscan Fast Inverted confocal microscope and contrast-adjusted in Photoshop or Fiji. For whole Matrigel dome images, scans were performed using a 10x objective on an inverted IX83 Olympus microscopy with a motorized XY-stage (Prior) and tiled images automatically generated by CellSens (Olympus).

## Immunohistochemistry

Tissue sections were deparaffinized with xylene and rehydrated with decreasing concentrations of ethanol (100%, 95%, 80%), then washed with ddH20. Antigen retrieval was performed with slides submerged in 10 mM citrate buffer (pH 6.0) and heated in a steamer for 90°C for 20min. Slides were cooled to room temperature and incubated with 6% H2O2 in methanol for 30 min. Following washing in 0.1% PBS-T (Phosphate-buffered saline, 0.1% Tween 20), slides were incubated in Avidin blocking solution for 15min, following by subsequent blocking in Biotin blocking solution for 15min (Vector Laboratories, SP-2001). Following washing in 0.1% PBS-T, slides were then incubated with serum-free Protein Block (Abcam, ab156024) for 10min. Sections were incubated with primary antibodies (cytokeratin-19 (Abcam, ab9221) and Ki67 (550609, BD Biosciences) diluted 1:250 in PBS-T overnight in a humidified chamber at 4°C. Next, slides were washed with PBS-T and incubated with secondary antibody (Biotinylated Goat Anti-Rabbit or Mouse IgG, Vector Biolabs BA- 1000 and BA-9200) for 30min, washed, and then incubated with avidin/biotin-based peroxidase (Vector Laboratories, Vectastain Elite ABC HRP Kit, PK-6100) for an additional 30min. Following washes in PBT-T, sections were incubated with DAB substrate (Vector Laboratories, SK-4100) for ∼5min. Slides were washed with ddH20 and then counterstained with hematoxylin for 1 min, thoroughly rinsed with H2O, and incubated in 0.1% sodium bicarbonate in H2O for 5 mins. Slides were then dehydrated with increasing concentrations of ethanol, cleared with xylene and mounted with Vectamount Permanent Mounting Medium (Vector Laboratories, H-5000). Images were captured on an IX83 inverted microscope (Olympus) using a UC90 color CCD camera (Olympus).

### RNA extraction, cDNA synthesis and quantitative PCR

Total RNA was extracted with the Sigma GenElute total mammalian RNA miniprep kit following the manufacturer’s instruction and using the supplementary Sigma DNase digestion. RNA quality and concentration were determined using a Nanodrop ND-1000 Spectrophotometer. Total RNA was reverse-transcribed with the iScript cDNA synthesis kit (Bio-Rad) following the manufacturer’s instructions. Quantitative PCR was performed using the iQ SYBR Green Supermix (Bio-Rad, 1708882) on a CFX96 Touch Real-Time PCR Detection System (Bio-Rad). Gene expression was determined based on a ΔCT method normalized to actin. The expression level of C19MC miRNAs hsa-miR-516b-5p, hsa-miR-517a-3p, hsa-miR- 525-5p and reference gene RNU6 was quantified by using the MystiCq® MicroRNA® Quantitation System (Millipore Sigma) which includes miRNAs Poly(A) tailing, cDNA synthesis (MIRRT) and miRNA qPCR (MIRRM00). MicroRNAs were first polyadenylated in a poly(A) polymerase reaction, then reverse transcribed using an oligo-dT adapter primer according to the manufacturer’s protocol (Sigma-Aldrich, MIRRT). Individual microRNAs (microRNA-516b-5p, microRNA-517a-3p and microRNA-525-5p) were quantified in real-time SYBR^®^ Green RT-qPCR reactions with the specific MystiCq microRNA qPCR Assay Primer (MIRAP00475, MIRAP00477 and MIRAP00512) and the MystiCq Universal PCR Primer (MIRUP). Cq reading were normalized to the RNU6 internal reference by the 2^-ΔCq^ method.

### RNAseq analysis

For RNAseq analysis, RNA was isolated from organoids as described above. Purified Total RNA was verified by Thermo scientific Nanodrop one. The libraries were prepared by the Duke Center for Genomic and Computational Biology (GCB) using the Tru-Seq stranded total RNA prep kit (Illumina). Sequencing was performed on the NovaSeq 6000 by using 150-bp paired-end sequencing. The reads were aligned to the human reference genome (GRCh38) using QIAGEN CLC Genomics (v20). DESeq2 [49] was used to normalized count data and to do the differential gene expression analysis with a significance cutoff of 0.01 and a fold change cutoff of log_2_(2). Heatmaps and hierarchical clustering was performed in R using pheatmap package in R and were based on RPKM (reads per kilobase million). Files associated with RNA-seq studies have been deposited into Sequence Read Archive under accession number SRA. Data from PHT cells (accession SRP067137 and SRP109039) [10, 11, 50]. Principal component analyses were performed using pcaExplorer in R [51] or GraphPad Prism. Volcano plots were generated using the EnhancedVolcano package in R [52] or in Graphpad Prism.

### Luminex assays

All sample processing was performed in duplicates and each experiment was performed with at least three biological replicates. Luminex assays were performed using the following kits according to the manufacturer’s instructions: Bio-Plex Pro Human Inflammation Panel 1, 37-Plex kit (171AL001M; Bio-Rad) Bio-Plex Human Chemokine Panel, 40-plex (171AK99MR2; Bio-Rad), Bio-Plex Pro Human Cytokine Screening Panel, 48-Plex kit (12007283; Bio-Rad), hCG Human ProcartaPlex Simplex Kit (EPX010-12388-901) and CA125 (Mucin-16) Human ProcartaPlex Simplex Kit (EPX010-12437-901). Plates were read on MAGPIX (EMD Millipore) or Bio-Plex 200 (Bio-Rad) Luminex machines and analyzed using xPONENT (EMD Millipore) or Bio-Plex (Biorad) software.

### IFN-λ activity reporter assay

For IFN-λ activity detection from CM, HEK-Blue IFN-λ reporter cells were used according to the manufacture’s protocol. Briefly, IFN-λ reporter cells were plated into a 96-well plate, then CM was added and incubated at 37 °C for 24 hr. Supernatants (20 µl) were collected and then transferred into a 96-well plate and activity measured using QUANTI-Blue Solution (InvivoGen, rep-qbs) after incubating at 37 °C for 2 hr using a spectrophotometer at 655 nm. Each CM preparation was run in triplicate.

### Poly(I:C) treatment

For Poly (I:C) treatment, TOs or DOs were incubated with 10 µg/mL Poly (I:C) diluted in organoid complete growth media for ∼24 h, then supernatant was collected for Luminex-based multianalyte profiling or RNA isolated as described for qRT-PCR or RNASeq.

### Statistics and reproducibility

All experiments reported in this study have been reproduced using independent samples (tissues and organoids) from multiple donors. All statistical analyses were performed using Prism software (GraphPad Software). Data are presented as mean ± SD, unless otherwise stated. Statistical significance was determined as described in the figure legends. Parametric tests were applied when data were distributed normally based on D’Agostino-Pearson analyses; otherwise, nonparametric tests were applied. P values <0.05 were considered statistically significant, with specific P values noted in the figure legends.

## Author contributions

LY designed and performed experiments and participated in manuscript review and writing, ECS designed and performed experiments and participated in manuscript review and writing, CO performed experiments and participated in manuscript review and writing, JG provided placental tissue and participated in manuscript review and writing, CM performed tissue dissections and participated in manuscript review and writing, SP designed experiments and participated in manuscript review and writing, and CC designed and performed experiments, participated in manuscript review and writing, and secured funding.

## Acknowledgements

We thank Jon Boyle (University of Pittsburgh) for careful review of the manuscript and Alexandra Wells for assistance with bioinformatics. This project was supported by NIH AI145828 (C.B.C.).

---

**Supplemental Figure 1.**
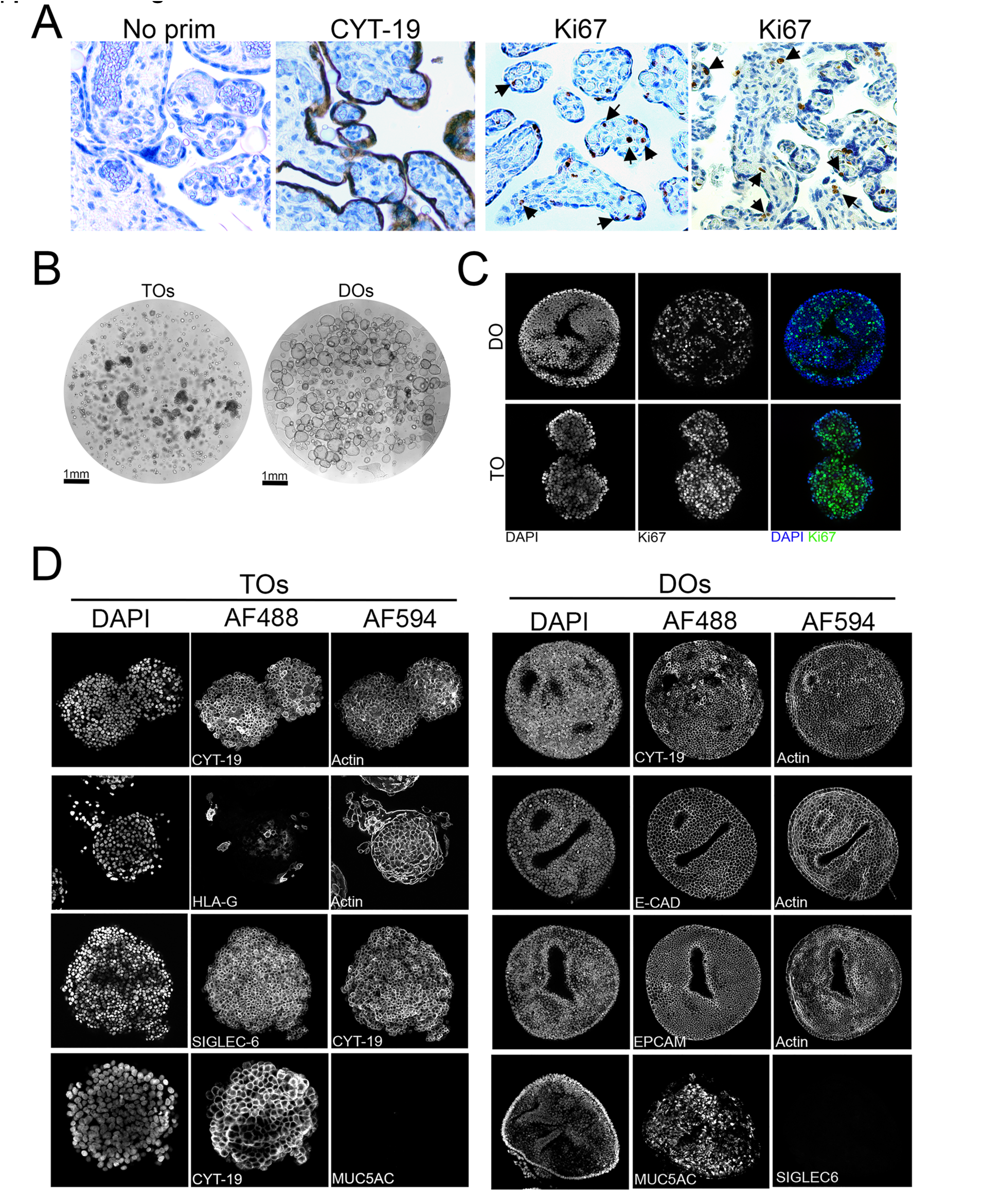
Characterization of trophoblast and decidua organoids. **(A)** Immunohistochemistry in human full-term placental villi tissue used to generate trophoblast organoids. Shown are no primary controls (left) and representative images of tissue immunostained for cytokeratin-19 (CYT-19) or Ki-67 as a marker of cell proliferation. Black arrows denote nuclei positive for Ki-67. **(B)** Brightfield images of Matrigel containing TOs (left) or DOs (right). Tiled images were captured using a motorized xy-stage at 10x magnification. Scale shown at bottom left. **(C)** Immunostaining for Ki67 (green) in DOs (top) or TOs (bottom). DAPI-stained nuclei are shown in blue. **(D)** Black and white images of single channel confocal micrographs used to generate the merged images shown in Figure 1C and D.

**Supplemental Figure 2.**
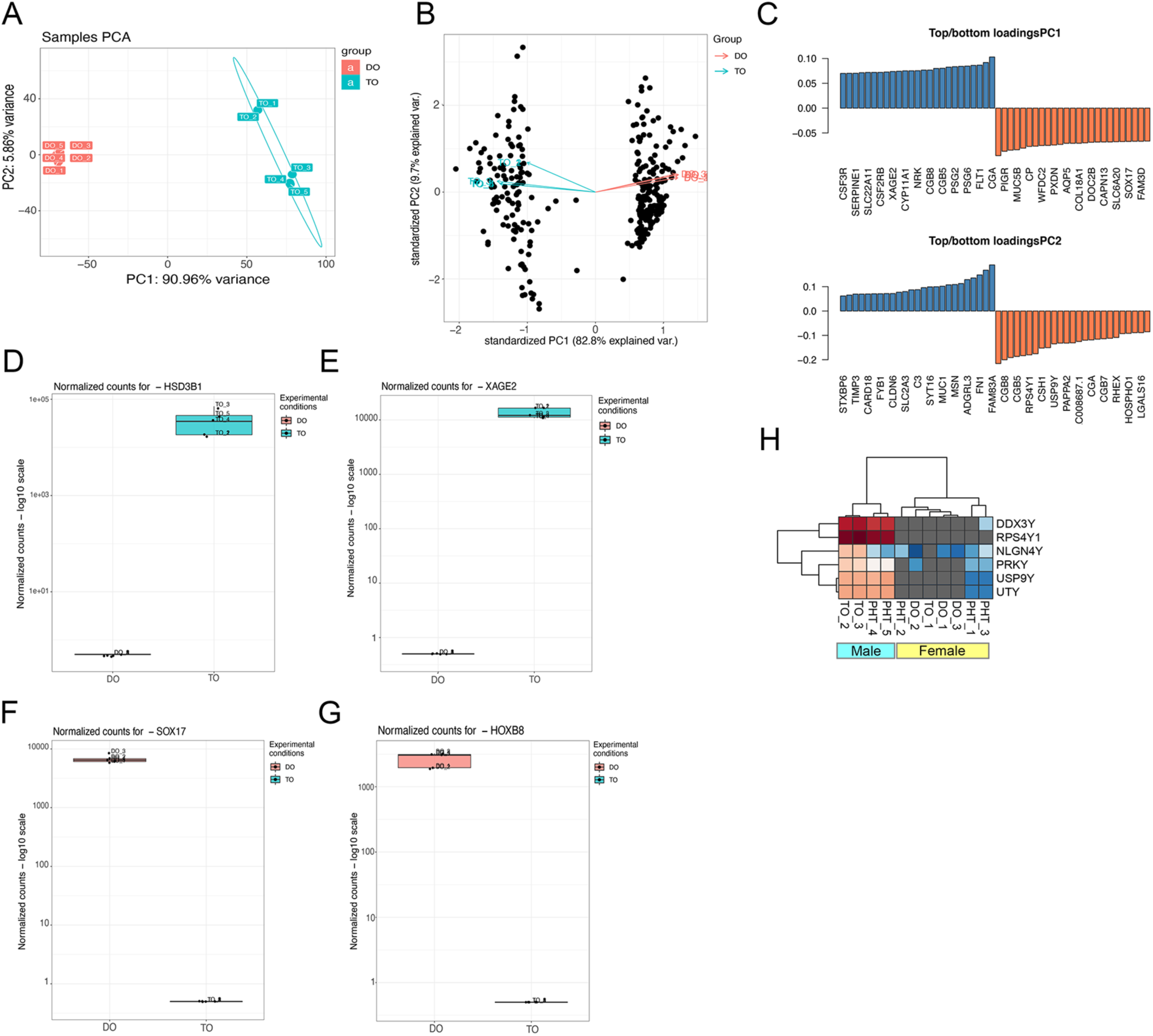
RNASeq of TOs and DOs. **(A, B)** Principal component analysis based on transcriptional profiles from DOs (orange) or TOs (blue**). (C)** Transcripts on the first and second principal components (PC1, PC2) with the greatest variance. Blue indicates enrichment in TOs and orange indicates enrichment in DOs. **(D-G)** Gene-specific enrichment of select transcripts in TOs (blue) and DOs (orange). Shown are HSD3B1 (D), XAGE2 (E), SOX17 (F), and HOXB8 (G). Five individual samples are shown as unique datapoints. **(H)** Heatmap (based on log2 RPKM values) of representative Y-linked transcripts in TOs, DOs, or primary human trophoblasts (PHT). Red indicates high levels of expression, blue indicates low levels of expression, and grey indicates no reads detected. Hierarchical clustering is shown at top.

**Supplemental Figure 3.**
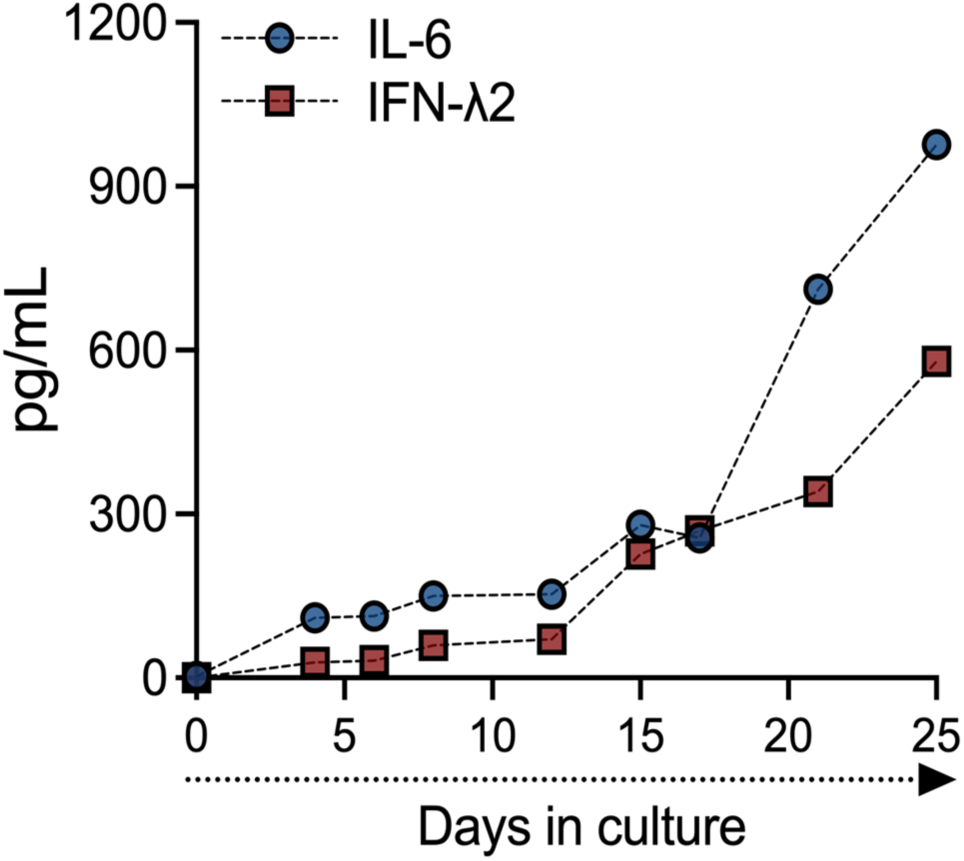
IL-6 and IFN-λ2 release in TOs over time. Concentrations of IL-6 (blue) and IFN-λ2 (red) in CM generated from a single TO line throughout the first 25 days of culture.

**Supplemental Figure 4.**
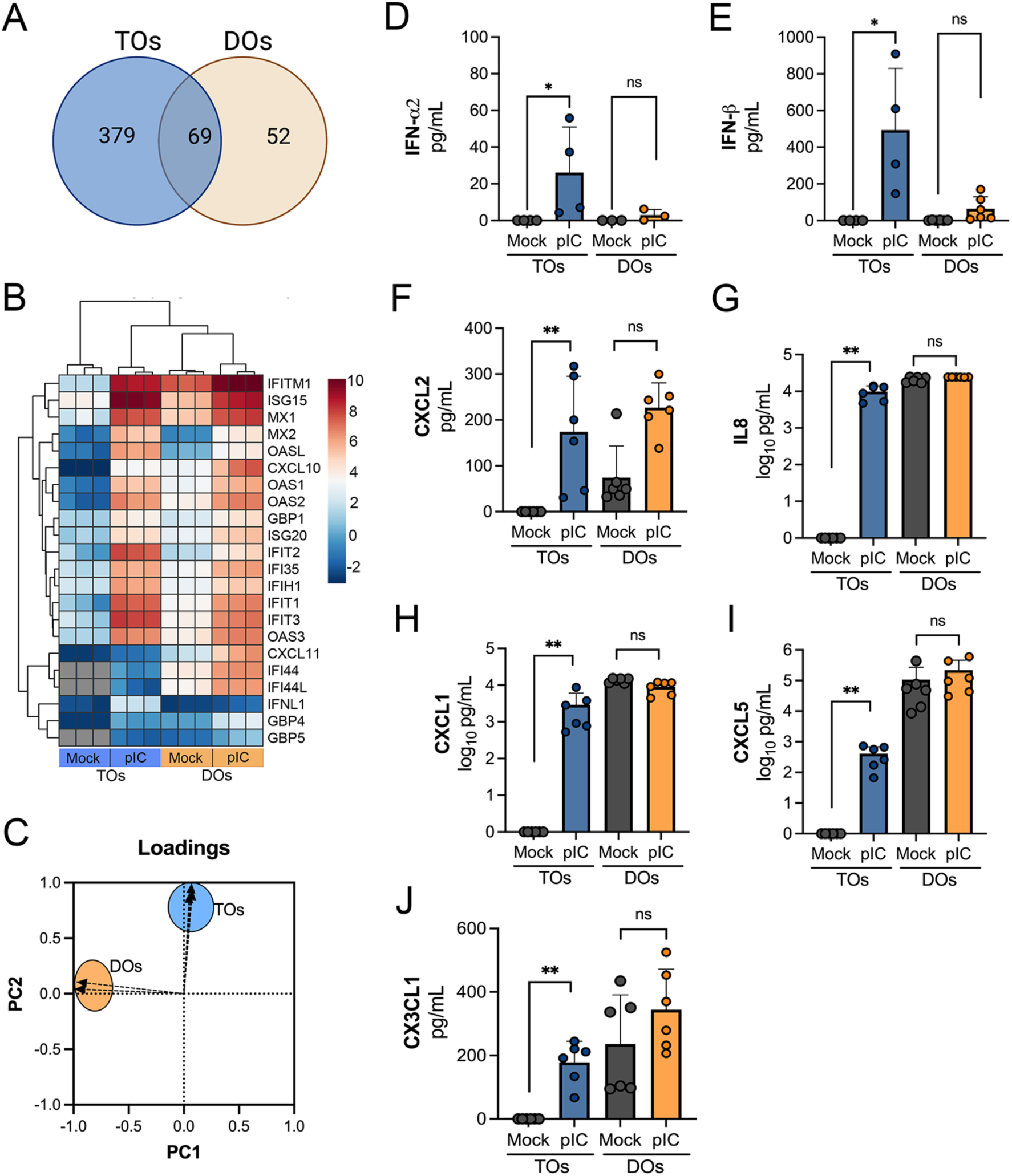
Factors induced by poly I:C treatment of TOs and DOs. **(A)** Venn diagram denoting unique and shared differentially expressed transcripts in TOs (blue circle) or DOs (orange circle) treated with poly I:C. **(B)** Heatmap (based on log2 RPKM values) of transcripts induced by poly I:C treatment of both TOs (blue box, bottom) or DOs (orange box, bottom). Key at right. Red indicates high levels of expression, blue indicates low levels of expression, and grey indicates no reads detected. Hierarchical clustering is on the left. **(C)** Loadings of principal component analysis of Luminex-based multianalyte profiling of poly I:C treated TOs (blue circle) and DOs (orange circle). **(D-J).** Levels of IFN-α2 (D), IFNβ (E), CXCL2 (F), IL-8 (G), CXCL1 (H), CXCL5 (I), and CX3CL1 (J) in CM isolated from poly I:C-treated TOs (blue) or DOs (orange) or in mock-treated controls (grey). Each symbol represents individual CM preparations. Significance was determined using a Mann-Whitney U test. ** P < 0.01, * P < 0.05; ns, not significant.

**Supplemental Figure 5.**
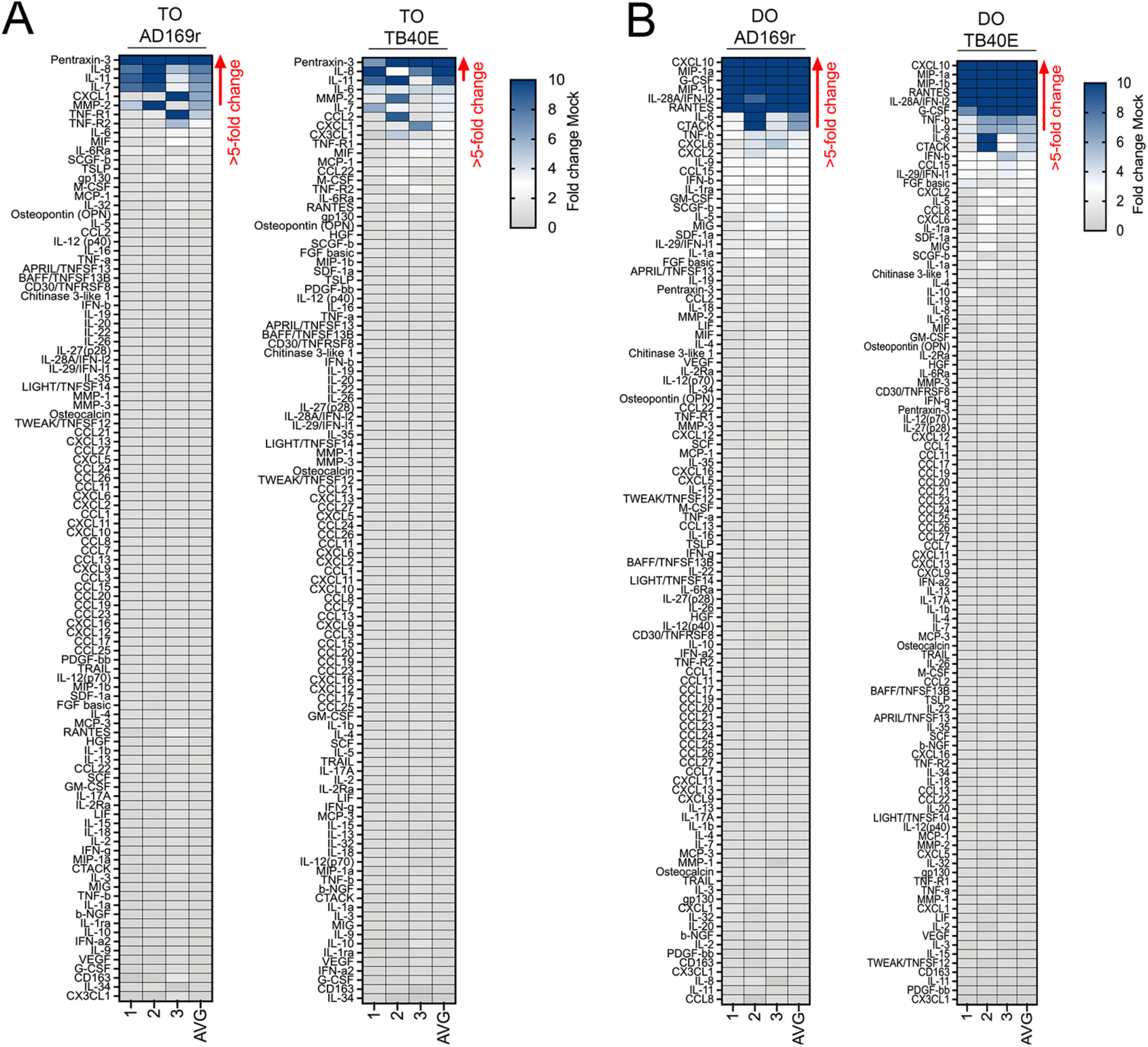
Differential innate immune responses in TOs and DOs infected with HCMV. **(A, B)** Heatmaps demonstrating the induction of factors at left (shown as fold change from mock-treated controls) from TOs (A) and DOs (B) infected with AD169r (left) or TB40E (right) strains of HCMV for ∼48hrs as assessed by Luminex-based multianalyte profiling. AVG denotes the average change in concentration of factors over conditioned medium (CM) isolated from three individual preparations. Dark blue denotes significantly induced factors compared with untreated controls. Gray or white denotes little to no change (scale at top right). The red arrow demonstrates factors with induction greater than five-fold change observed in the average of separate experiments.

